# Inverse and Proportional *trans* Modulation of Gene Expression in Human Aneuploidies

**DOI:** 10.1101/2023.09.21.558906

**Authors:** Shuai Zhang, Ruixue Wang, Ludan Zhang, James A. Birchler, Lin Sun

## Abstract

Genomic imbalance in aneuploidy is often detrimental to organisms. To gain insight into the molecular basis of aneuploidies in humans, we analyzed transcriptome data from several autosomal and sex chromosome aneuploidies. The results showed that in human aneuploid cells, genes located on unvaried chromosomes are inversely or proportionally *trans*-modulated, while a subset of genes on the varied chromosomes are compensated. Less genome-wide modulation is found for sex chromosome aneuploidy compared with autosomal aneuploidy due to X inactivation and the retention of dosage sensitive regulators on both sex chromosomes to limit the effective dosage change. We also found that lncRNA and mRNA can have different responses to aneuploidy. Furthermore, we analyzed the relationship between dosage-sensitive transcription factors and their targets, which illustrated the modulations and indicates genomic imbalance is related to stoichiometric changes of components of gene regulatory complexes.

## Introduction

The concept of gene balance has been known for a hundred years in that partial genomic dosage changes are highly detrimental on the phenotypic level (Birchler and Veitia, 2021). This principle is upheld in multicellular eukaryotes including humans. Molecular studies of aneuploidy have been varied and have yielded different interpretations as to the underlying basis of the detrimental phenotypes. One such idea is that the varied region exhibits a gene dosage effect for the encoded genes. However, it is now realized that many types of regulatory genes are dosage sensitive and that they will modulate target loci across the genome indicating a more global aspect to the basis of aneuploid syndromes (Birchler and Veitia, 2021). Although gene expression levels are found to be proportional to their copy numbers in some aneuploidy experiments, there is strong evidence that in many cases, aneuploidy can also perturb the expression of genes on other chromosomes, based on the identity of the varied chromosome and the context (Kojima and Cimini, 2019).

Several observations indicate the impact of dosage sensitive regulatory genes across the genome. First, retained duplicates from the two whole genome duplications in the vertebrate lineage are those with dosage sensitive action on multiple targets and which cause syndromes when varied in dosage (Ionita-Laza et al., 2009; Makino and McLysaght, 2010; Singh et al., 2012; Session et al., 2016; Mottes et al., 2021; Zhang D et al., 2021). A complementary pattern of underrepresentation of dosage sensitive regulators in small scale duplications occurs (Maere et al., 2005; Blomme et al., 2006; Freeling, 2009; Gout and Lynch, 2015; Tasdighian et al., 2017; Mottes et al., 2021; Zhang D et al., 2021). This complementary pattern appears related to that of multicomponent regulatory genes in parallel to aneuploid effects (Birchler et al., 2005; Birchler and Veitia, 2012, 2021) and confirmed by a similar complementary pattern involving Protein-Protein Interactions (Defoort et al., 2019). MicroRNA genes, which quantitatively modulate the levels of many transcription factors, are also held in duplicate following whole genome duplications in vertebrates (Berthelot et al., 2014; Desvignes et al., 2021; Peterson et al., 2022). Further, the importance of the effect of dosage sensitive regulatory genes within a two-fold range is illustrated by the preferential retention of them in three independent sex chromosomal systems in vertebrates (Bellott et al., 2014, 2017; Cortez et al., 2014; Bellott and Page, 2021). Whereas most genes can be deleted to a single copy in one or the other sex, depending on the type of sex chromosomal system, the dosage sensitive regulators are maintained on both sex chromosomes. It has been suggested that members of multisubunit complexes are compensated on the protein level by degradation (Dephoure et al., 2014; Hwang et al., 2021), but the evolutionary results noted above would not be realized if this were universally the case. Also, the fact that most transcription factors are dosage sensitive (Seidman and Seidman, 2002; Veitia, 2002) would suggest that most target genes would be modulated across the genome.

The behavior of dosage sensitive genes involved in multicomponent interactions shows parallels to aneuploidy/ploidy genomic balance phenomena as previously noted (Birchler et al., 2001, 2005; Birchler and Veitia, 2012, 2021). Single dosage sensitive genes can mimic the effect of aneuploidy on a larger scale and are involved with various regulatory functions (Rabinow et al., 1991; Birchler et al., 2001; Zhang S et al., 2021b). Taken together with current knowledge of gene regulatory networks, these considerations suggest a need to examine data from human aneuploids for genome wide modulations of gene expression.

The across-genome modulation of aneuploidy has been described in detail in various species such as maize, *Drosophila*, and *Arabidopsis* (Birchler and Veitia, 2021). Early studies in maize have found that dosage changes of large chromosome segments produce *trans*-acting effects on the remainder of the genome, which may be positive or negative, but it is more common that the regulation of expression levels is negatively correlated with copy number changes (i.e. inverse dosage effect) (Birchler, 1979; Birchler and Newton, 1981; Guo and Birchler, 1994). This effect is thought to act as a dominant negative impact of differentially varying the individual components of regulatory complexes (Birchler and Veitia, 2021). Across organisms in which the global effects of aneuploidy have been studies, the most common response closely approximates an inverse correlation with the magnitude of the aneuploidy but can exhibit a range. Here we collectively refer to the down modulation in hyperploidy as an “inverse effect”.

At the same time, the expression of some genes encoded by the varied chromosomes did not change with the gene dosage; in other words, they were compensated (Birchler, 1981; Birchler and Newton, 1981; Guo and Birchler, 1994). In *Drosophila*, both autosomal and sex chromosome aneuploidies can affect the expression levels of other chromosomes, while the transcription of the varied chromosome is closer to normal diploid levels (Devlin et al., 1982, 1988; Birchler et al., 1990; Sun et al., 2013a,b,c; Zhang S et al., 2021a). Recent whole-genome studies of maize and *Arabidopsis* aneuploid series have also demonstrated the prevalence of the inverse dosage effect and dosage compensation (Hou et al., 2018; Johnson et al., 2020; Shi et al., 2021; Yang et al., 2021). The mechanism of dosage compensation in aneuploidy was regarded as the simultaneous negative genome-wide effect counteracting the positive gene dosage effect, and was verified by subdivision of chromosome arms (Birchler, 1981; Birchler et al., 1990).

In order to examine the genome-wide modulation patterns of gene expression in human aneuploidy, and to investigate whether the previously found *trans*-acting effects and dosage compensation exist in human, we performed an analysis of RNA sequencing data from public databases for human autosomal aneuploidies (trisomy 13, 18, and 21) (Letourneau et al., 2014; Sullivan et al., 2016; Hwang et al., 2021) and sex chromosome aneuploidies (XO and XXY) (Zhang X et al., 2020; Astro et al., 2022). We found that both inverse and proportional *trans* modulation are present in human aneuploid cells, and dosage compensation also occurs for a subset of genes on the varied chromosomes. We also found a stronger response of lncRNAs than mRNAs to aneuploid variation. Sex chromosome aneuploidy shows many fewer genome wide modulations despite the fact that the X chromosome is of substantial size, which illustrates the effectiveness of X inactivation to mitigate the potentially harmful effects of dosage sensitive regulatory genes. In addition, the relationship of dosage-sensitive transcription factors and their influence on targets was analyzed. Together, these studies indicate that human aneuploidy causes extensive genome-wide modulation of gene expression in a similar fashion as occurs in other species.

## Results

### Dosage compensation and inverse dosage effects in human aneuploid cells

To investigate the global impact of aneuploid variation on the human genome, we first analyzed a set of RNA-seq data from trisomies 13, 18, and 21 (Hwang et al., 2021). This dataset is particularly amenable because a single cell type (fibroblasts) from multiple trisomies was studied. By calculating the ratio of gene expression in aneuploidy to the corresponding normal diploid and generating ratio distributions of all expressed genes (filtering out lowly expressed genes) on the varied chromosomes (*cis*) and unvaried chromosomes (*trans*) (Figure 1A-F), it is possible to reveal the overall trend and subtle impacts of gene expression modulation.

**Figure 1.**
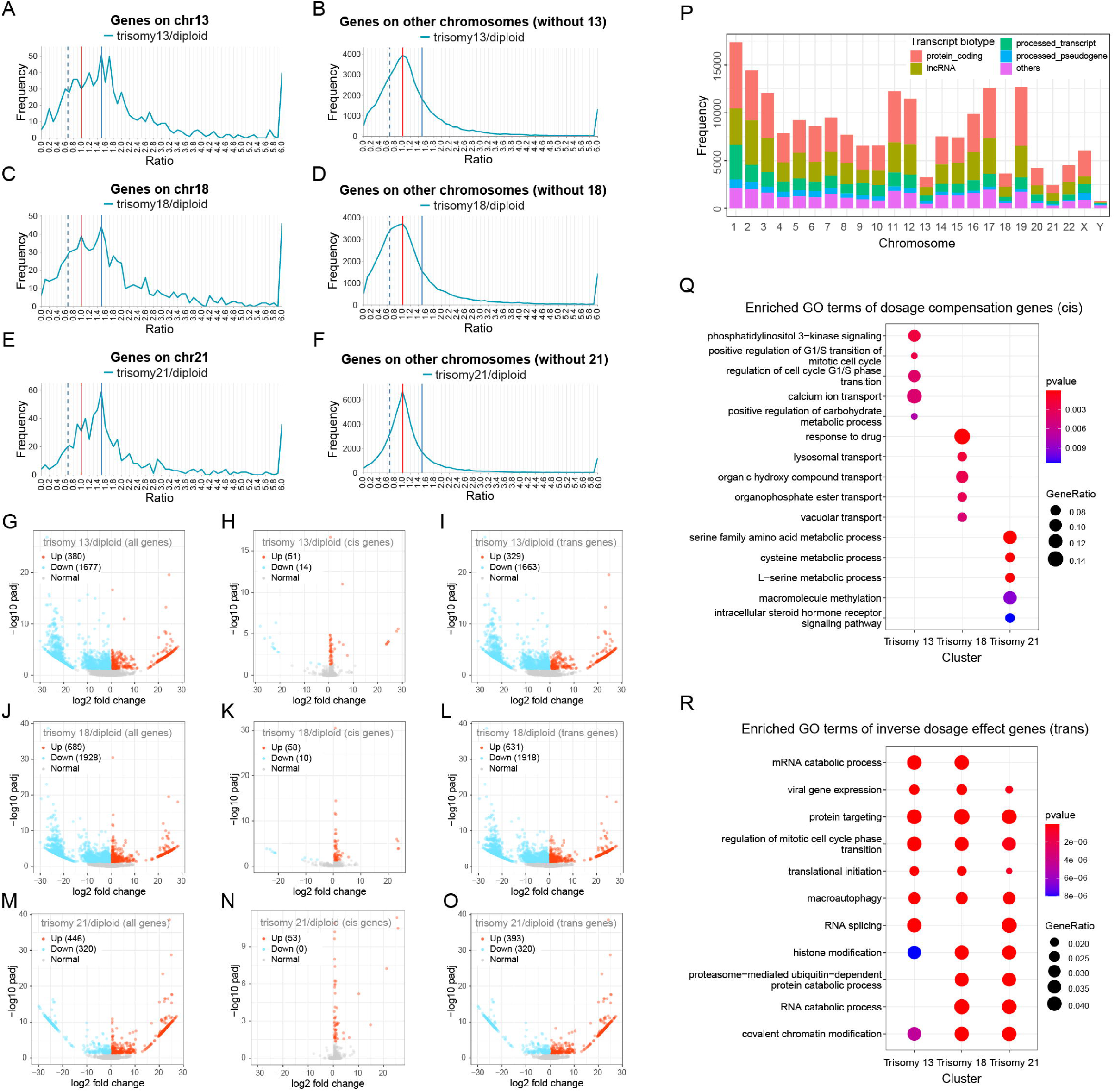
Global expression analysis of human autosomal aneuploidy. (**A**-**F**) Ratio distributions of gene expression in human autosomal aneuploidies compared with normal individuals. The ratio distributions of *cis* and *trans* genes in trisomy 13 (**A** and **B**), trisomy 18 (**C** and **D**), and trisomy 21 (**E** and **F**) are shown, respectively. The X-axis indicates the ratio of gene expression, and the Y-axis indicates the frequency of the ratios that fall into each bin of 0.1. The red vertical line represents the ratio of 1.00, and the blue vertical dashed and solid lines indicate 0.67 and 1.5, respectively. Sample size: trisomy 13, n = 2; trisomy 18, n = 2; trisomy 21, n = 7; diploid, n = 6. (**G**-**O**) Volcano plots of differentially expressed genes (DEGs) in human autosomal aneuploidies. (**G**-**I**) Volcano plots of DEGs in trisomy 13 compared with normal diploid. (**J**-**L**) Volcano plots of DEGs in trisomy 18 compared with normal diploid. (**M**-**O**) Volcano plots of DEGs in trisomy 21 compared with normal diploid. Genes located on all chromosomes (**G**, **J** and **M**), *cis* genes (**H**, **K** and **N**), and *trans* genes (**I**, **L** and **O**) are shown respectively. DEGs are defined as adjusted p-value < 0.05. Significantly up-regulated genes are represented by red dots and significantly down-regulated genes are represented by blue dots. The legend in the upper left corner shows the number of up-regulated and down-regulated transcripts in each plot. (**P**) The numbers and types of transcripts on each human chromosome. (**Q**) Functional enrichment analysis of *cis* genes showing typical dosage compensation (ratios between 0.9 and 1.1). (**R**) Functional enrichment analysis of *trans* genes showing typical inverse dosage effect (ratios between 0.57 and 0.77).

In the ratio distribution plots, a ratio of 1.0 will be obtained if the expression level of a gene does not change between the aneuploid and the diploid (Figure 1A-F). The Kolmogorov-Smirnov test (K-S test) showed that all distributions do not fit to a normal distribution with a mean of 1.0 (Appendix 1—table 12), indicating that the *cis* and *trans* genes are not normally distributed in the three autosomal aneuploidies. For genes located on the triplicated chromosomes, multiple prominent peaks are observed in all distributions (Figure 1A,C,E), the highest one at 1.5, representing a proportional increase in gene expression (dosage effect), and the lower at 1.0, representing unchanged levels of expression (dosage compensation). There are also small peaks near the inverse level (0.67), indicating a reduction in expression in the trisomy even below the diploid level. These results suggest that in human aneuploid cells, a subset of *cis* genes is expressed in direct relationship to their copy number, while others show compensation and gradations between and beyond these levels. RNA-seq studies provide a relative but not absolute relationship of the mRNAs in the transcriptome (Loven et al., 2012). In this regard, the inverse effect cannot be due to *cis* gene dosage effects skewing the relative reads because the landscapes of the *cis* distributions are multidimensional as are the *trans* landscapes. If sequencing skewing in *cis* and *trans* were occurring, single peaks offset from 1.5 and 1.0, respectively, with no other peaks would be predicted but this was not found. Critically, the addition of the proportional increases to the total transcriptome from these chromosomes (∼ 0.4-0.6%) is not sufficiently large relative to the remainder of the genome to produce such a skew.

For genes on the unvaried chromosomes, the ratio distributions spread around 1.0 (Figure 1B,D,F). The main peak of 1.0 means that a proportion of *trans* genes are unaffected by the addition of a chromosome. However, these peaks are skewed to the left in all aneuploidies. Especially in trisomy 18 a shoulder peak is found below 1.0, which also contains a considerable number of genes, approximately at the ratio of 0.67, representing the negative effect of changes in chromosome numbers (inverse dosage effect). In addition, there are also some *trans* genes whose ratio increases, and the number of genes located near 1.5 (ratios between 1.4 and 1.6) is about half of those at 0.67 (ratios between 0.57 and 0.77), indicating a positive *trans*-acting effect. Therefore, dosage changes on one chromosome can have widespread effects throughout the human genome, not only affecting its own gene expression, but also regulating other parts of the genome. The *trans*-acting modulations have both proportional and negative effects, but the inverse dosage effect is more predominant.

In order to determine whether the normalization method will affect the results of the ratio distributions, we used two additional normalization methods, TPM (Transcripts Per Million) and TMM (Trimmed Mean of M-values), to generate the ratio distributions (Figure 1—figure supplement 1; Figure 1—figure supplement 2). It is found that the major patterns are essentially the same using all three types of normalization. Therefore the normalization methods will not affect the analysis. In addition, we also generated ratio distributions for each human chromosome in trisomies 13, 18, and 21 (Figure 1—figure supplement 3; Figure 1—figure supplement 4; Figure 1—figure supplement 5). The results show that the ratio distributions of most *trans* chromosomes are similar to the overall *trans* distribution, suggesting that the dosage effects broadly affect the expression of genes on all chromosomes.

It has been suggested that the regulation in aneuploidy may also correlate with gene expression levels (Stamoulis et al., 2019). Therefore, we divided all genes into low, medium, and high expression groups, and the number of transcripts in each group is similar (Figure 1—figure supplement 6A-F). We found that the genes in the low and medium level groups tend to show dosage compensation in *cis* and inverse modulation in *trans* (Figure 1—figure supplement 6A-F, left two panels) compared with the high-expression group due to their general left-shift peaks (K-S test p-values < 0.05, Appendix 1—table 12). Genes in the high-expression group tend to show a dosage effect more consistently in *cis*, while genes in *trans* are less modulated (Figure 1—figure supplement 6A-F, rightmost panels).

In addition to the ratio distributions, we also determined the differentially expressed genes (DEGs) in these autosomal aneuploidies (Figure 1G-O; Appendix 1—table 1). For trisomy 13, the down-regulated genes (1677) are much greater in number than up-regulated genes (380), and a similar trend of more down regulated genes (1928) than upregulated genes (689) was found for trisomy 18 (Figure 1G,J). For trisomy 21 with fewer total DEGs, slightly fewer down-regulated genes are found than up-regulated genes (Figure 1M). Since the total number of *cis* genes is small, the global changes of the genome mainly come from the changes of *trans* genes. As seen in the volcano plots, in trisomies 13 and 18, the expression of a large number of *trans* genes shows significant changes, most of which are down-regulated (Figure 1I,L). This result is consistent with the peaks of inverse dosage effect in the ratio distribution plots and the general left skew of the *trans* distributions (Figure 1B,D). Most *cis* DEGs are up-regulated, but there are some *cis* genes whose expression is significantly down-regulated (Figure 1H,K). In addition, there is no significant difference in the expression of most *cis* genes (∼ 90%), meaning that the addition of one chromosome does not result in significant upregulation of most genes on that chromosome, implying dosage compensation.

After dividing the DEGs into low, medium and high expression groups (Appendix 1—table 2; Appendix 1—table 3; Appendix 1—table 4), we found that in trisomies 13 and 18, the number of significantly down-regulated transcripts in each group is greater than that of up-regulated transcripts. Although a small fraction of the highly-expressed *cis* genes are significantly up-regulated, which show a dosage effect, the majority of the low and medium expression *cis* genes have no significant difference in expression levels. For *trans* genes in trisomies 13 and 18, the total number of DEGs is small, but the number of down-regulated genes (∼ 200) was five times that of the up-regulated genes. And the number of down-regulated genes with medium and high expression are similar, approximately 700 to 800, higher than the corresponding number of up-regulated genes.

Notably, we included all the transcripts in this analysis, not only protein-coding genes but also non-coding RNAs. Statistical analyses on the types and numbers of genes on the 24 human chromosomes show that long non-coding RNAs account for a high proportion on each chromosome, which together with protein-coding genes constitute the majority of transcription types (Figure 1P). Thus, the effects observed above are equally effective for both protein-coding genes and non-coding RNAs. Furthermore, chromosomes 13, 18, and 21 are the three autosomes containing the lowest number of transcripts among all human chromosomes (Figure 1P), so the trisomy of these chromosomes may have a relatively small global impact compared to others, which is consistent with the relative survival of various human aneuploidies (Orr et al., 2015). Based on the number of transcripts from trisomic chromosomes, we found that the *cis*-distributed dosage compensation peak and the *trans*-distributed inverse effect peak were more prevalent with trisomy 18 (Figure 1C,D). In contrast, chromosome 21, with the lowest number of transcripts, had lesser genome-wide effects (Figure 1F). We hypothesize that the modulation of gene expression in aneuploidy may be related to the size of the varied chromosomes, which would change correspondingly the number of dosage sensitive regulatory genes. A larger genomic segment of imbalance results in stronger global regulation, which is more likely to induce dosage compensation in *cis* and an inverse dosage effect in *trans*.

Functional enrichment analysis was performed for genes that exhibited typical dosage compensation (ratios between 0.9 and 1.1) and *trans*-acting inverse dosage effects (ratios between 0.57 and 0.77; Figure 1Q,R). The results showed that the *cis* compensated genes had various functions and had no similarities among the three aneuploidies (Figure 1Q). In addition, there is no similarity in the functions of *cis* genes showing a dosage effect (Figure 1—figure supplement 7A). The functions of the more finely divided *cis* genes appear to be more dispersed (Figure 1—figure supplement 7B). The functions of *trans* genes that exhibit an inverse dosage effect are mainly focused on RNA processing and metabolism, protein targeting, autophagy, and chromatin modification, all of which are important for cell regulation (Figure 1R). The *trans* genes affected by the positive dosage effect (ratios between 1.4 and 1.6) are also enriched for RNA metabolism, splicing, localization, and chromatin regulation (Figure 1—figure supplement 7). The unchanged *trans* genes (ratios between 0.9 and 1.1) have some different functions, including protein metabolism, vesicle transportation *et cetera* (Figure 1—figure supplement 7D). In general, aneuploid variation leads to changes in expression of many functional types of genes, and the functions of dosage compensated *cis* gene are heterogeneous.

### Relationship between dosage compensation and the inverse dosage effect

Among these human aneuploidies, we found that in trisomy 18 a large subset of *cis* genes was closer to dosage compensation, and the *trans* modulations were primarily inversely affected (Figure 1C,D), which were validated by differential gene expression analysis (Figure 1J-R). This observation indicates that the *trans* genes showing an inverse dosage effect and the *cis* genes predisposed to compensation are likely affected by related mechanisms as previously found (Birchler et al., 1990; Hou et al., 2018; Shi et al., 2021; Yang et al., 2021; Zhang S et al., 2021a). To investigate whether the dosage compensated genes on each chromosome were refractory to modulation, we generated *trans* distributions for the expression of *cis* compensated genes (DC genes; ratios between 0.9 and 1.1) identified in each trisomy for an effect by the other trisomies (Figure 2A-F). In other words, we tested whether dosage compensated genes in each trisomy could be inversely or otherwise modulated by the other trisomies. The normality tests show that four of the six *trans* distributions of these genes fit a normal distribution (Appendix 1—table 12), indicating the stability of the expression of DC genes. The distribution of DC genes shows high and sharp peaks at the ratio around 1.0 (Figure 2A-F, red line), indicating that the expression levels of a subset of DC genes remain unchanged by any *trans*-acting effects from trisomies of other chromosomes, although this does not exclude the possibility that still other trisomies might modulate them. However, all the distributions show an obvious secondary peak at a ratio approximating 0.67, representing the inverse dosage effect, which is higher and sharper than that of all genes on the respective chromosome (Figure 2A-F). This result indicates that at least some dosage compensated genes from one trisomy can be modulated by the other trisomies. By tabulating the number of overlapping *cis* DC genes and *trans* inversely affected genes (IDE genes; ratios between 0.57 and 0.77) on chromosomes 13, 18, or 21, we found that the *cis* DC genes from each trisomy are significantly enriched in the IDE genes of other trisomies (Fisher’s exact test p-value < 0.05; Figure 2G-I). In a reciprocal analysis, we determined the *cis* distributions of genes on one chromosome that showed an inverse dosage effect (IDE) in one of the other trisomies (Figure 2J-L). Significantly different from the *cis* distributions of all genes on these chromosomes (K-S test p-value < 0.05, Appendix 1—table 12), the distributions of IDE genes are more concentrated. Peaks less than 1.5 are present in these *cis* distributions, suggesting that the expression of some IDE genes can be inversely modulated when their copy numbers are increased. Therefore, we believe that there is a relationship between the *cis* gene-by-gene dosage compensation and the *trans* gene-by-gene inverse dosage effect in accordance with results from other organisms (Hou et al., 2018; Shi et al., 2021; Yang et al., 2021).

**Figure 2.**
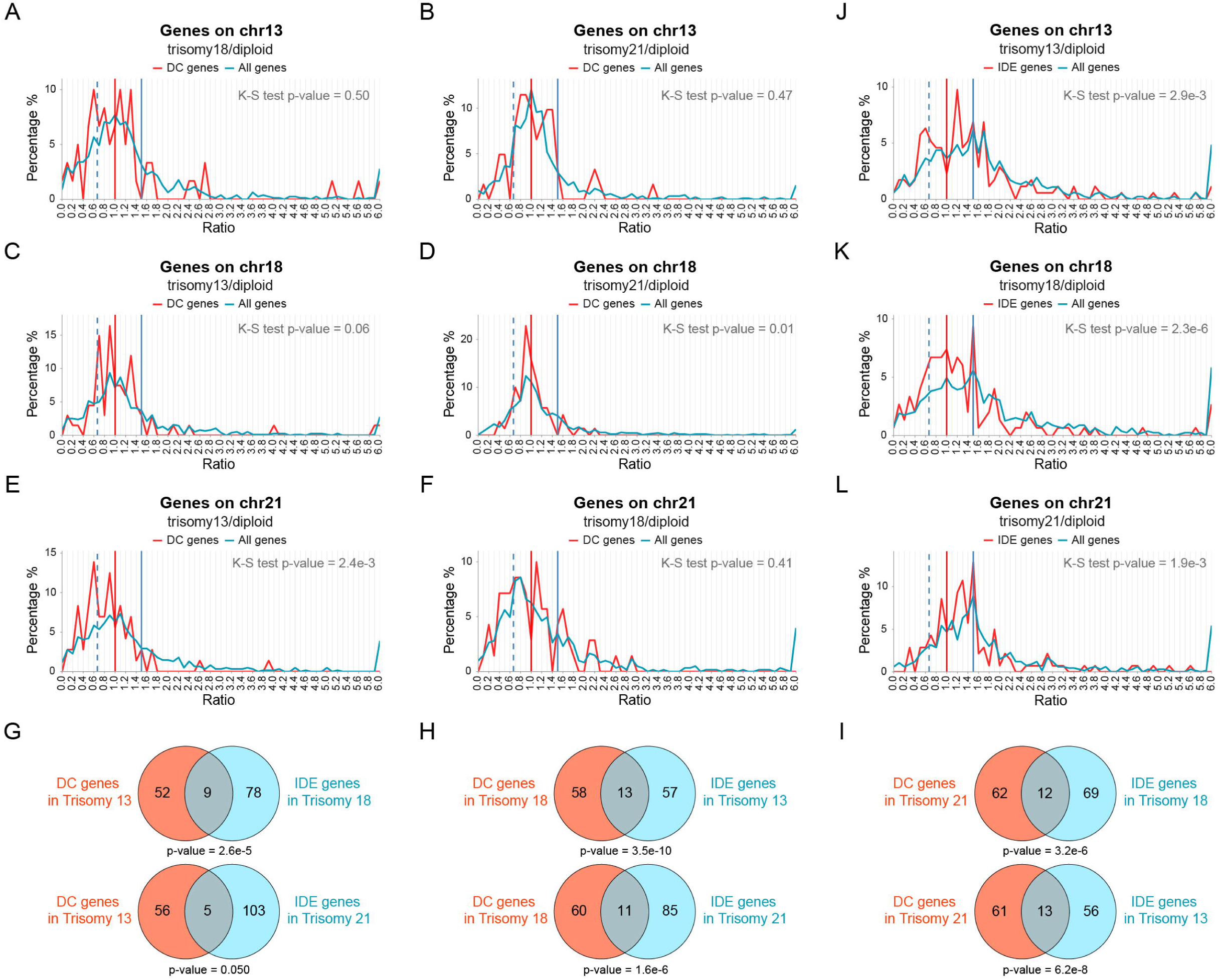
The relationship between dosage compensation and an inverse dosage effect. (**A**-**F**) Ratio distributions of the expression of dosage compensated (DC) genes (red lines) and all genes on this chromosome (blue lines) for the other two human autosomal aneuploidies compared with normal individuals. The *trans* distributions of *cis* DC genes (ratios between 0.9 and 1.1) identified in trisomy 13 (**A** and **B**), trisomy 18 (**C** and **D**), and trisomy 21 (**E** and **F**) are shown for the other two aneuploidies, respectively. (**G**-**I**) Venn diagrams of DC genes and inverse dosage effect (IDE) genes (ratios between 0.57 and 0.77). The numbers of *cis* DC genes identified in one aneuploidy and *trans* IDE genes identified in other two aneuploidies that are located on chromosomes 13 (**G**), 18 (**H**), and 21 (**I**) are shown. Fisher’s exact test p-value are listed below each plot. (**J**-**L**) Ratio distributions of the expression of IDE genes (red lines) and all genes on this chromosome (blue lines) in three human aneuploid types compared with normal individuals. The *cis* distributions of *trans* IDE genes identified and combined in the other two aneuploidies are shown in trisomy 13 (**J**), trisomy 18 (**K**), and trisomy 21 (**L**), respectively.

### Aneuploid effects analyzed by age, sex, and genetic background

To examine further dosage compensation and *trans*-acting dosage effects in human aneuploidy, two other trisomy 21 sequencing datasets of human fibroblasts were analyzed (Figure 3). The first dataset includes a pair of monozygotic twins with discordant chromosomes 21 with a consistent genetic background, and a group of unrelated trisomy 21 and normal individuals (Letourneau et al., 2014). According to the ratio distributions, the main *cis* peak in trisomy 21 of the twins is located near 1.0, and the secondary peak in *cis* is at the ratio of 1.5 (Figure 3A), indicating that most genes on chromosome 21 are compensated, and some genes also show a proportional gene dosage effect without compensation. The distribution of *trans* genes is skewed (does not fit a normal distribution, Appendix 1—table 12) and the expression levels of most genes are down-regulated, forming a single peak at approximately 0.8, which is negatively correlated with the dosage changes (Figure 3B).

**Figure 3.**
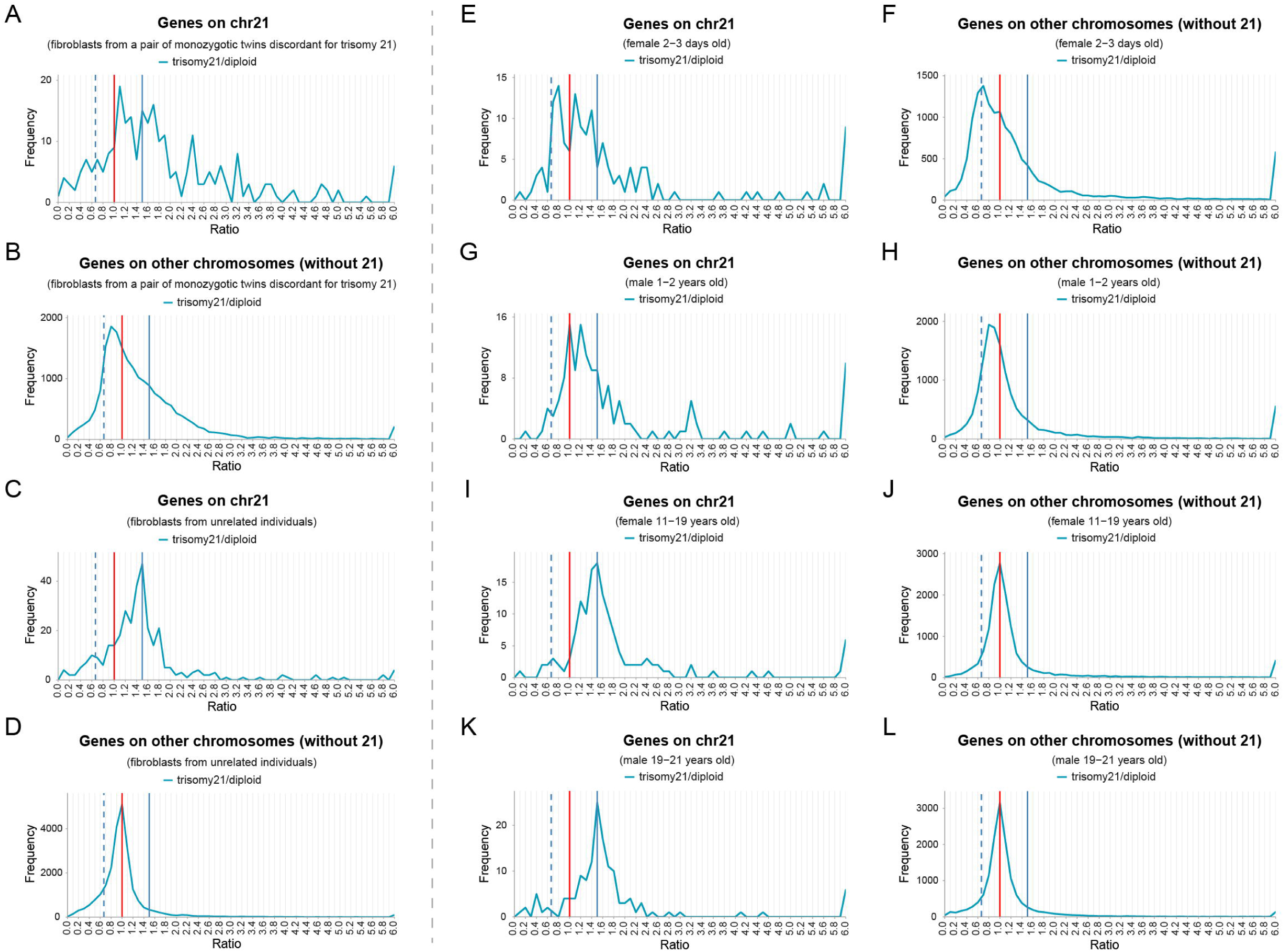
Ratio distributions of gene expression in two human trisomy 21 datasets. (**A**-**B**) Ratio distributions of gene expression of fibroblasts collected from a pair of trisomy 21 and normal discordant monozygotic twins. Sample size: trisomy 21, n = 4; diploid, n = 4. (**C**-**D**) Ratio distributions of fibroblast gene expression in a group of unrelated trisomy 21 and normal individuals. Sample size: trisomy 21, n = 8; diploid, n = 8. (**E**-**L**) Ratio distributions of gene expression of fibroblasts collected from trisomy 21 and normal individuals of the same sex and similar ages, which are divided into four groups, 2-3 days old females (**E**-**F**), 1-2 years old males (**G**-**H**), 11-19 years old females (**I**-**J**), and 19-21 years old males (**K**-**L**). Sample size: 2-3 d female trisomy, n = 2; 2-3 d female diploid, n = 2; 1-2 y male trisomy, n = 4; 1-2 y male diploid, n = 2; 11-19 y male trisomy, n = 4; 11-19 y male diploid, n = 2; 19-21 y male trisomy, n = 2; 19-21 y male diploid, n = 4.

In the comparison of samples from unrelated individuals, the main *cis* peak is located at 1.5, which represents the dosage effect, and the secondary peak is between 1.0 and 1.5, representing some degree of compensation (Figure 3C). The ratio distribution of *trans* genes does not deviate from 1.0, indicating that the *trans*-acting effects are slight. Thus, dosage compensation and *trans*-acting dosage effects are not detected to the same degree in this dataset in which differences in genetic background are present.

Another set of data includes sequencing results of fibroblasts from individuals of different ages and sexes (Sullivan et al., 2016). We assigned these samples to four groups of matched ages and sexes, and analyzed the distribution of their gene expression ratios, respectively (Figure 3E-L). The analysis found that, in the comparison of two younger groups, *cis* genes show obvious compensation, and *trans* genes tend to show an inverse dosage effect, with distributions skewed to the left and do not fit a normal distribution (Appendix 1—table 12; Figure 3E-H). However, in the older groups, the *cis* genes show more proportional dosage effects, while the *trans* genes remain unchanged (K-S test p-values < 0.05 between the young and old groups, Appendix 1—table 12; Figure 3I-L). The limited number of samples, imperfect age matching, and individual variation make conclusions about the influence of these categories difficult. In other systems any one aneuploidy can modulate the same gene differently in different cell types (Cooper and Birchler, 2001) and this is the case in human as well (Liu et al., 2023). Nevertheless, our analysis of these trisomy 21 datasets demonstrates that all of them exhibit gene-by-gene proportional dosage effects, dosage compensation, and inverse dosage effects, as a general rule, similarly to the other comparisons described above.

### LncRNA and mRNA can have different responses to aneuploidy

LncRNAs have been implicated in a variety of regulatory processes (Statello et al., 2021). Because there is little research on the effect of lncRNA expression levels in aneuploidy, we generated separate ratio distributions for mRNA and lncRNA in three autosomal aneuploidies (Figure 4A-F). As for protein-coding genes, the dosage effect peaks (1.5) are higher than compensation peaks (1.0) in *cis* distributions, and the shoulder peaks of the inverse dosage effect (∼ 0.67) in *trans* are less than the unmodulated peak (Figure 4A-F, red line). In contrast, the ratio distributions of lncRNA are shifted to the left in trisomy 18 (K-S test p-value = 9.1e-5 in *cis* and p-value < 2.2e-16 in *trans*), indicating down-regulation of expression (Figure 4A-F, blue line). In the *cis* distributions of trisomy 18 and 21, peaks representing dosage compensation at 0.9-1.1 are more obvious for lncRNAs than for mRNAs (Figure 4C,E; Figure 4G, gray boxes). Among these *trans* distributions, the peak approximating 0.67 is also more obvious for lncRNA than mRNA (K-S test p-values < 0.05, Appendix 1—table 12; Figure 4B,D,F). The proportion of lncRNA with an inverse modulation on unvaried chromosomes is also greater than that of mRNA in trisomies 18 and 21 (Figure 4G, blue boxes).

**Figure 4.**
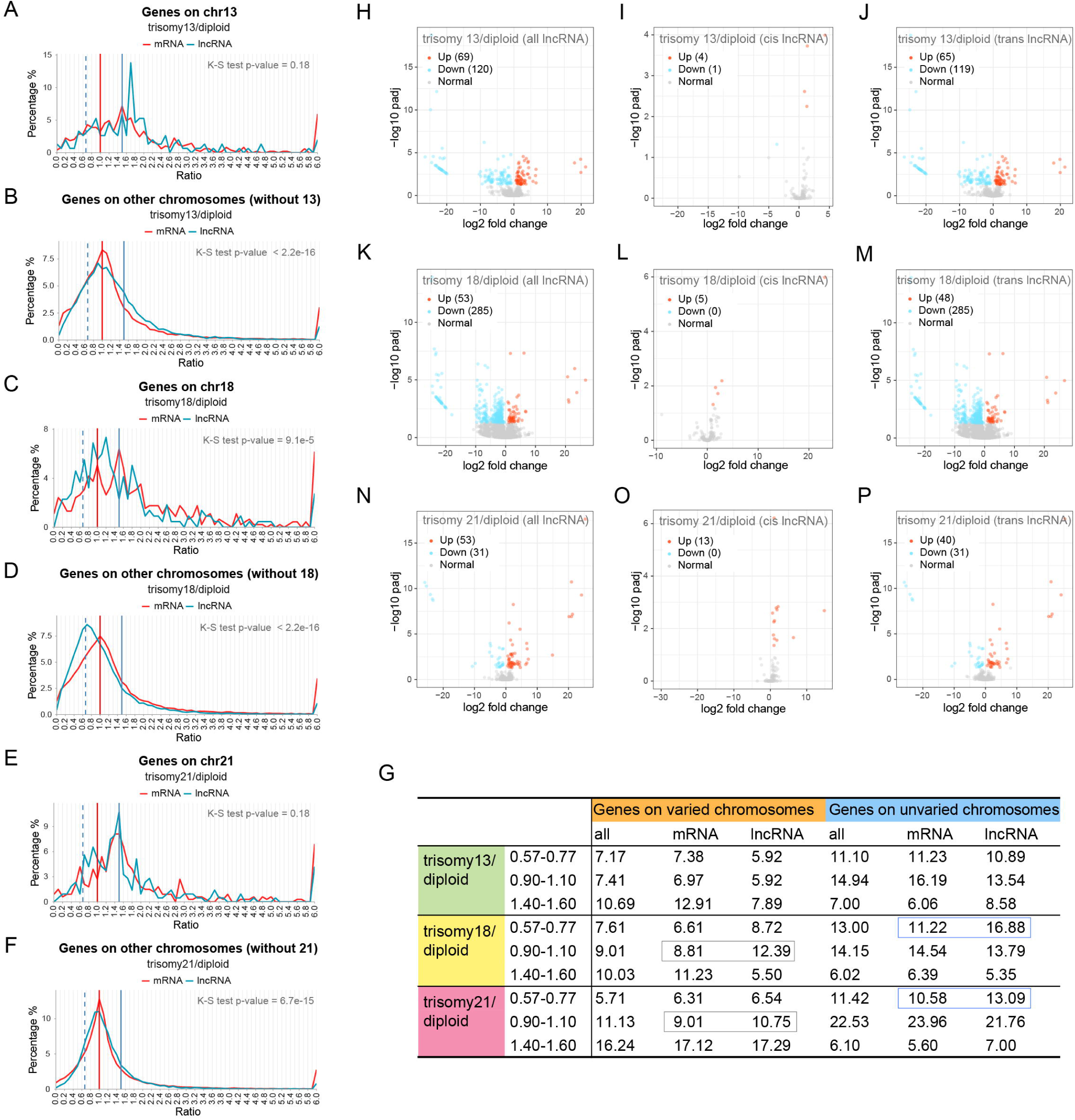
Comparison of lncRNA and mRNA in aneuploidy. (**A**-**F**) Ratio distributions of lncRNA and mRNA transcription in human autosomal aneuploidies compared with normal individuals. The ratio distributions of *cis* and *trans* lncRNA (blue lines) and mRNA (red lines) in trisomy 13 (**A** and **B**), trisomy 18 (**C** and **D**), and trisomy 21 (**E** and **F**) are shown, respectively. (**G**) The table shows the percentage of lncRNA and mRNA on the varied and unvaried chromosomes within specified ratio ranges. The blue boxes indicate the typical inverse dosage effect of *trans* genes, and the gray boxes indicate dosage compensation. (**H**-**P**) Volcano plots of differentially expressed lncRNA (DE-lncRNA) in human autosomal aneuploidies. (**H**-**J**) Volcano plots of DE-lncRNA in trisomy 13 compared with normal diploid. (**K**-**M**) Volcano plots of DE-lncRNA in trisomy 18 compared with normal diploid. (**N**-**P**) Volcano plots of DE-lncRNA in trisomy 21 compared with normal diploid. Genes located on all chromosomes (**H**, **K** and **N**), *cis* lncRNA (**I**, **L** and **O**), and *trans* lncRNA (**J**, **M** and **P**) are shown respectively. DE-lncRNAs are defined as adjusted p-value < 0.05. Significantly up-regulated lncRNAs are represented by red dots and significantly down-regulated lncRNAs are represented by blue dots. The legend in the upper left corner shows the number of up-regulated and down-regulated transcripts in each plot.

We also used TPM and TMM to generate ratio distributions for lncRNA and mRNA (Figure 4—figure supplement 1; Figure 4—figure supplement 2). In the results of these two additional normalization methods, it was also observed that the distributions of lncRNA were more left skewed than that of mRNA. We also generated ratio distributions for each chromosome in the human complement (Figure 4—figure supplement 3; Figure 4—figure supplement 4; Figure 4—figure supplement 5), and found that the expression of lncRNA is more down-regulated than that of mRNA on most *trans* chromosomes in accordance with the original analysis.

Similarly, we also generated volcano plots for significantly differentially expressed mRNA and lncRNA, respectively (Figure 4—figure supplement 6; Figure 4H-P). Similar to the above, for the protein-coding genes of trisomies 13 and 18, the significantly down-regulated *trans* lncRNA transcripts are greater in number than the up-regulated *trans* transcripts, and most of the *cis* transcripts are not significantly up-regulated; instead, they are compensated to some extent (Figure 4—figure supplement 6A-F). The DEG analysis for lncRNAs supports the results of the ratio distributions (Figure 4H-P; Figure 2A-F).

In addition, we also divided mRNA and lncRNA into low, medium and high groups according to their expression levels for analysis (Figure 4—figure supplement 7 and Figure 4—figure supplement 8). Among the three aneuploidies, the mRNAs with low and medium expression mainly exhibit an inverse correlation with dosage changes, while high-expression mRNAs are more likely to show a dosage effect in *cis* and be unchanged in *trans* (K-S test p-values < 0.05 for low and medium groups compared with high groups, respectively, Appendix 1—table 12; Figure 4—figure supplement 7A,C,E; Figure 4—figure supplement 8A,C,E). However, the differences in the distributions of lncRNAs among the three expression groups on either varied or unvaried chromosomes are not as obvious as mRNAs (Appendix 1—table 12; Figure 4—figure supplement 7B,D,F; Figure 4—figure supplement 8B,D,F). For trisomy 18 with the largest aneuploid effects, the *cis* distributions of low, medium, and high expression lncRNAs tend to compensate, and the peak of *trans* distribution approximates an inverse relationship (Figure 4—figure supplement 7D; Figure 4—figure supplement 8D). Furthermore, all the *trans* distributions and most of the *cis* distributions of mRNAs and lncRNAs are significantly different among the different expression levels (Appendix 1—table 12; Figure 4—figure supplement 7; Figure 4—figure supplement 8). The results of differential expression analysis of mRNA and lncRNA of different expression levels (Appendix 1—table 2; Appendix 1—table 3; Appendix 1—table 4) are similar to that of all transcripts, as described above.

Through these analyses, we found that lncRNA and mRNA can have different responses to aneuploidy. lncRNAs can be more sensitive than mRNA to dosage modulation, and *cis* lncRNAs are more often compensated, while accordingly *trans* lncRNAs are more susceptible to an inverse modulation.

### The effects of sex chromosome aneuploidy are minimal due to compensation mechanisms

In addition to the autosomal aneuploidy described above, we also analyzed the changes of global gene expression in a group of human sex chromosome aneuploidies (Zhang X et al., 2020) (Figure 5A-F). In normal male and female, the expression of most of the genes on the X chromosome are similar although the copy numbers are different, with the peak of ratio distribution at 1.0 (Figure 5A, gray line). The distribution on the autosomes is concentrated at 1.0 (Figure 5B, gray line). This observation suggests that most of the genes are transcribed similarly in normal male and female, both on the autosomes and sex chromosomes. Further, after a reduction of X chromosome dosage in females in Turner syndrome, the peak of the distribution is still at 1.0 (Figure 5A, red line). Similarly, the XXY (Figure 5A, blue line) and XY (Figure 5A, gray line) genotypes also have nearly overlapping distributions, although their X chromosome numbers are different. The ratio distributions of genes located on the autosomes are more concentrated and almost overlap (Figure 5B). Thus, to some extent, the dosage changes in the X and Y chromosomes do not appear to have much effect on the entire gene expression global pattern due to the existence of X inactivation. This conclusion is also verified by the results of differential expression analysis, in which the total number of DEGs in each group is much smaller than that of autosomal aneuploidy, and the numbers of up-regulated and down-regulated genes are similar (Appendix 1—table 5). Further, the addition of a Y chromosome in XXY compared with XX does not change the expression of most genes (Figure 5A, blue line; Appendix 1—table 5). The effective change in dosage involves only the small pseudoautosomal regions of the X and Y chromosomes, and genes that escape inactivation. Nevertheless, Zhang and colleagues (Zhang X et al., 2020) noted a small subset of autosomal genes that were modulated in both genotypes with the predominant effect being inverse. Analysis of mRNA and lncRNA revealed that both have similar responses to X chromosome aneuploidies as all types of genes, but the *cis* and *trans* distributions of lncRNA seem to be more dispersed (Appendix 1—table 12; Figure 5C-F). However, lncRNAs do not show a higher proportion of differentially expressed transcripts (Appendix 1—table 5).

**Figure 5.**
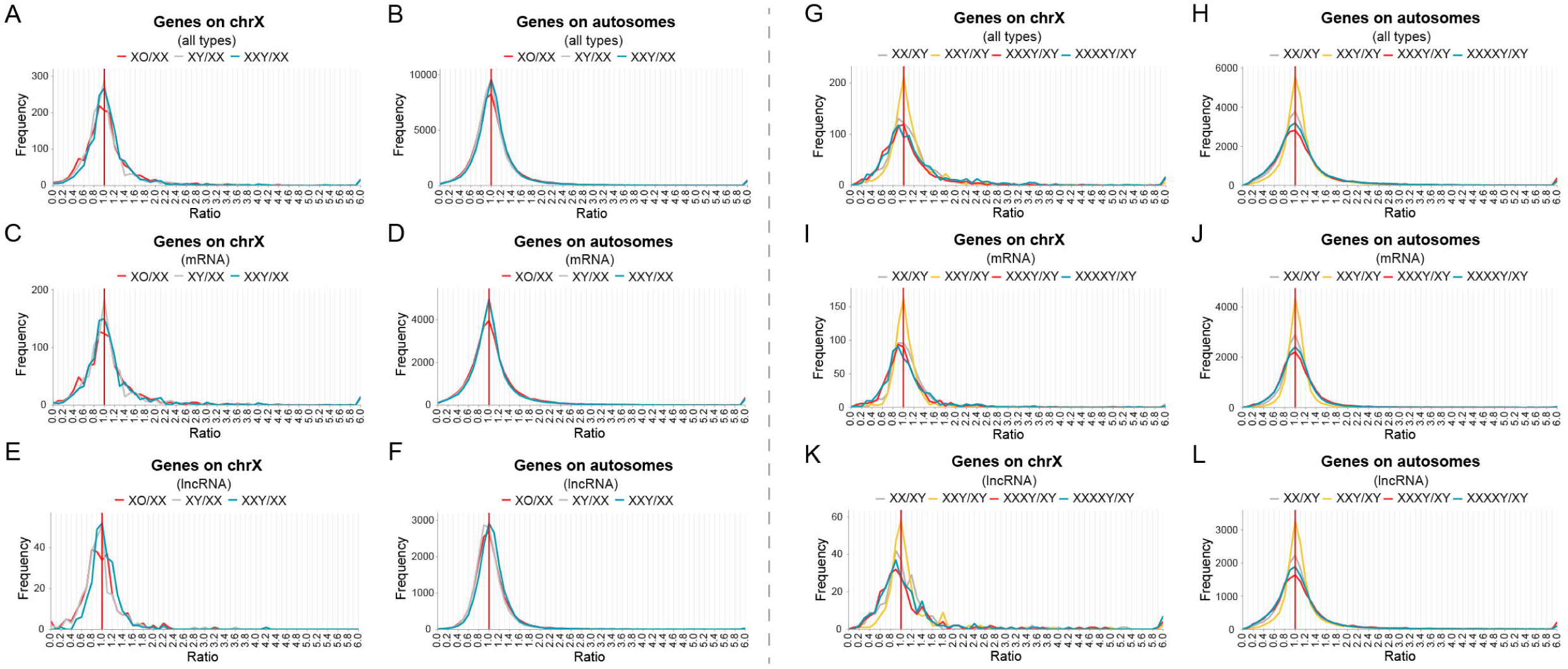
Global expression analysis of human sex chromosome aneuploidy. (**A**-**F**) Ratio distributions of gene expression in XO and XXY aneuploidies compared with normal individuals. The ratio distributions of all genes (**A** and **B**), protein-coding genes (**C** and **D**) and lncRNA (**E** and **F**) on the X chromosome and the autosomes are shown, respectively. Sample size: XO, XXY, and XX, n = 14; XY, n = 13. (**G**-**L**) Ratio distributions of gene expression in XXY, XXXY, and XXXXY aneuploidies compared with normal individuals. The ratio distributions of all genes (**G** and **H**), protein-coding genes (**I** and **J**) and lncRNA (**K** and **L**) on the X chromosome and the autosomes are shown, respectively. Sample size: XY, n = 13; XXY, n = 24; XXXY, n = 2; XXXXY, n = 4. The X-axis indicates the ratio of gene expression, and the Y-axis indicates the frequency of the ratios that fall into each bin of 0.1. The red vertical line represents the ratio of 1.00.

To further examine the effects of sex chromosome aneuploidy, we analyzed an additional set of sequencing data from Klinefelter syndrome (XXY) and high-grade X chromosome aneuploidies (XXXY and XXXXY) (Astro et al., 2022) (Figure 5G-L). Because of X inactivation the effective dosage change involves only the pseudoautosomal regions and those escaping inactivation. The results showed that although the number of X chromosomes increased from 2 to 4 copies in these aneuploidies, the global gene expression levels of both the X chromosome and autosomes did not shift obviously, and the ratio distribution peaks are all located near 1.0 (Figure 5G,H). For XXY, the proportions of down-regulated genes on both the X chromosome and the autosomes are lower than that of up-regulated genes (Appendix 1—table 6). However, with the further increase in X chromosome dosage, the ratio distributions of the X chromosome and the autosomes in high-grade aneuploidies are significantly different from that of XXY (K-S test p-values < 0.05 for all *cis* and *trans* distributions, Appendix 1—table 12), and the proportion of down-regulated genes is doubled (XXXY ∼ 35%, XXXXY ∼ 30%, ratio < 0.8; Appendix 1—table 6). The proportions of up-regulated genes on the X chromosome and the autosomes did not change so dramatically (∼ 20-30%, ratio > 1.25; Appendix 1—table 6). The results of *t*-tests further support the above statistics (Appendix 1—table 7). The number of significantly down-regulated genes increased in high-grade aneuploidies (XXXY and XXXXY) compared with XXY, and is more than the number of up-regulated genes (Appendix 1—table 7). Thus, with the increased degree of imbalance in sex chromosome aneuploidy, the magnitude of genome-wide inverse modulation is intensified. Nevertheless, the X inactivation mechanism weakens the disturbance of gene expression on the sex chromosome and autosomes, making the ratio distributions near a constant level. The results of mRNA and lncRNA were similar to the overall analysis (Figure 5I-L), and the proportions of up- and down-regulated lncRNAs are larger than the corresponding mRNAs (Appendix 1—table 6). These results are also consistent with previous conclusions (Astro et al., 2022).

### Dosage-sensitive transcription factors induce the inverse modulation of target genes

We hypothesized that the global effects of aneuploidy are due to dosage-dependent regulatory genes located on the varied chromosomes affecting target genes across all chromosomes. Previous studies have found that dosage-sensitive regulators usually involve multi-subunit complexes or components of multiple interactions, such as transcription factors, signal transduction components, and chromatin proteins (Birchler et al., 2001; Maere et al., 2005; Blomme et al., 2006; Makino and McLysaght, 2010; Tasdighian et al., 2017; Birchler and Veitia, 2021). Therefore, we studied the expression changes of transcription factors on varied chromosomes as well as their targets (Figure 6; Appendix 1—table 8-11). It is observed that the distributions of transcription factors located on the varied chromosomes have more spread peaks due to the small number involved (Appendix 1—table 12; Figure 6A,C,E).

**Figure 6.**
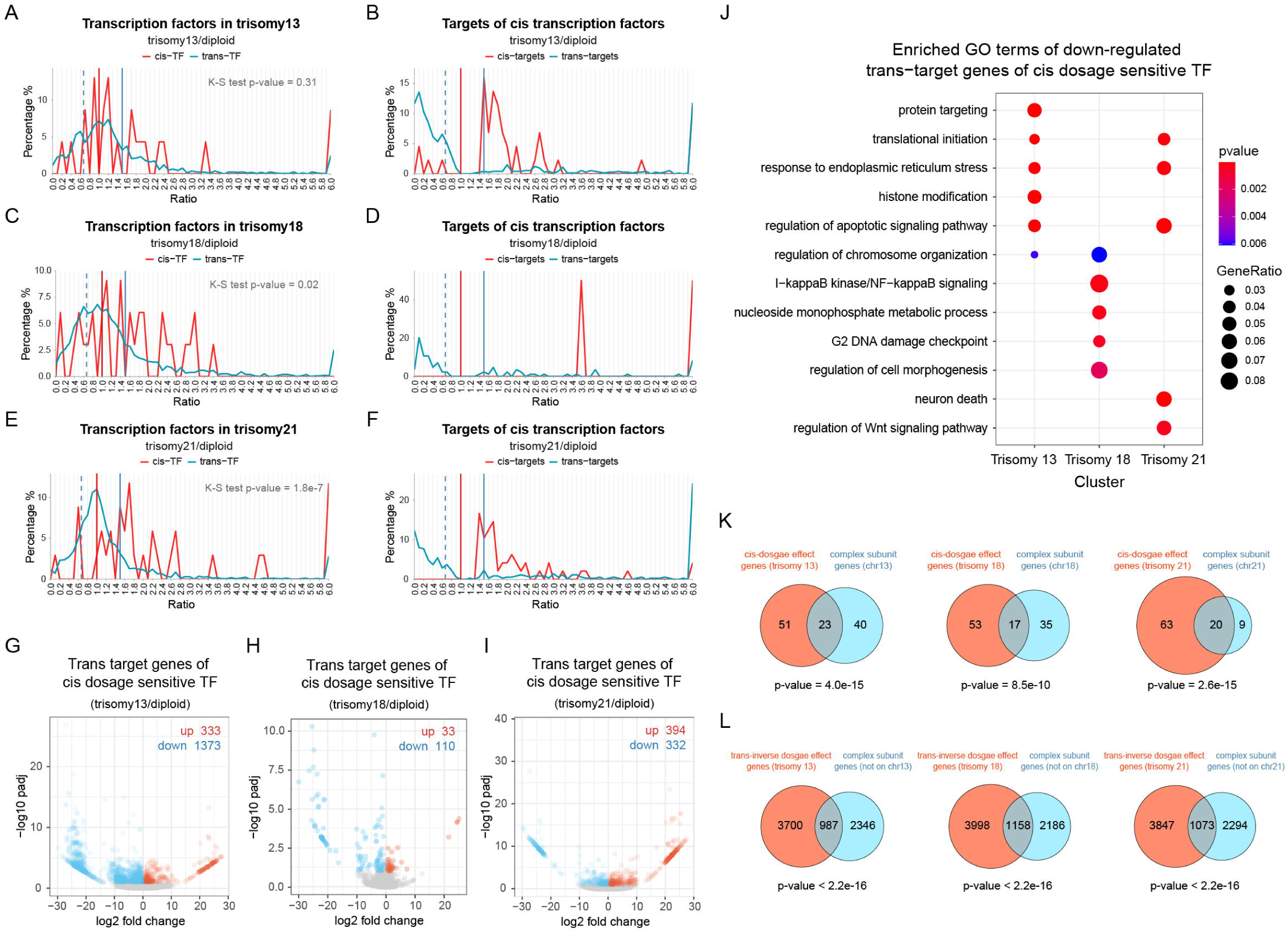
Expression of transcription factors and their target genes. (**A**-**F**) The ratio distributions of *cis* and *trans* transcription factors (TFs) (**A**, **C**, **E**) and the *cis* and *trans* differentially expressed target genes of *cis*-TFs (**B**, **D**, **F**) in human autosomal aneuploidies compared with normal individuals are shown, respectively. (**G**-**I**) Volcano plots show the expression of *trans* target genes of *cis* dosage-sensitive TFs in trisomy 13 (**G**), trisomy 18 (**H**), or trisomy 21 (**I**). The numbers of up-regulated and down-regulated genes are indicated in the top right corner of each plot. (**J**) Functional enrichment analysis of down-regulated *trans* target genes of *cis* dosage-sensitive TFs. Dosage sensitivity of transcription factors is defined as an increase in expression of more than 1.25-fold with the increasing copy numbers. A target gene is considered to be differentially expressed if the adjusted p-value is less than 0.1. (**E** and **F**) Venn diagrams show the overlap of macromolecular complex genes with *cis*-dosage effect genes (ratio between 1.4 and 1.6; **E**) and *trans*-inverse dosage effect genes (ratio between 0.57 and 0.77; **F**). The genes of macromolecular complex subunits were obtained from the core set in the CORUM database.

Next, we studied the *cis* and *trans* targets of transcription factors located on the varied chromosomes. In order to show the changing trend more clearly, we only generated ratio distributions for significantly differentially expressed target genes (Figure 6B,D,F). Most of the differentially expressed *cis* target genes have increased expression levels, but the total of gene numbers is small (Figure 6B,D,F, red line). On the contrary, most of the *trans* target DEGs are significantly down-regulated (Figure 6B,D,F, blue line), reflecting the *trans*-acting effects of *cis* transcription factors. To further examine this point, we analyzed all *trans* target genes of *cis* dosage-sensitive transcription factors, which we define as these with an expression more than 1.25-fold (Figure 6G-I). Trisomy 13 and trisomy 18 are observed to have more down-regulated *trans* target genes than up-regulated target genes, while trisomy 21 shows a similar number of up and down target genes. The specific numbers of target genes for each group of transcription factors are shown in Appendix 1—table 8-11. Furthermore, we also analyzed the functions of these down-regulated target DEGs, and found that there are few similarities among different aneuploidies (Figure 6J) indicating that the inverse effect is not specific to a particular functional category of genes.

By analyzing the distribution of genes contained in the CORUM database of human protein complexes, we found that the subunits of macromolecular complexes are significantly enriched in *cis*-genes with proportional dosage effects (ratios between 1.4 and 1.6, Fisher’s exact test p-value < 0.05; Figure 6K). Similarly, protein complex types are also significantly enriched in genes on the remaining chromosomes that are inversely affected (ratios between 0.57 and 0.77, Fisher’s exact test p-value < 0.05; Figure 6L). In addition, more than half of the dosage-sensitive transcription factors in each aneuploidy described above are confirmed to have interactors. Thus, at least at the RNA level these aneuploids will have stoichiometric changes for members of macromolecular complexes, which would likely change at least the initial translational stoichiometry of the complex members.

## Discussion

In this study, we analyzed the global gene expression in several human aneuploids and found that dosage compensation for subsets of genes on the varied chromosomes and *trans*-acting dosage effects for subsets of genes on the unvaried chromosomes are prevalent. We also examined the response of lncRNAs and the effect of sex chromosomes. Further, we analyzed the changes of gene expression regulatory networks throughout the genome with emphasis on dosage-sensitive transcription factors. These analyses provide new insights into human aneuploid effects on gene expression.

The concept of genetic imbalance has been widely reported for over a century, describing the phenomenon that changing the dosage of individual chromosomes or large chromosomal segments (aneuploidy) has more severe effects on the phenotype than changing a whole set of chromosomes in a genome (ploidy) (Birchler and Veitia, 2012, 2021). In human, only trisomies 13, 18, and 21 of the entire autosomal aneuploidies can survive to birth, while only trisomy 21 can survive to adulthood (Orr et al., 2015; Sanchez-Pavon et al., 2021). This fact may be related to the size of chromosomes or the number of genes contained, with chromosome 21 encoding the fewest genes and causing the least global disruption of gene expression.

The molecular basis of genomic imbalance in aneuploidy was initially thought to be due to the proportional dosage effects of gene expression produced by genes on varied chromosomes, which to some degree is correct, but genome-wide modulations are a critical component (Birchler and Veitia, 2012, 2021). Forty years ago, studies on enzyme and protein levels of maize aneuploid series revealed that the expression of some genes located on unvaried chromosomes were inversely regulated and coincidentally dosage compensation was found for a subset of genes on the varied chromosomes (Birchler, 1979, 1981; Birchler and Newton, 1981; Guo and Birchler, 1994). Since then, the inverse dosage effect and dosage compensation of aneuploidy have been reported in different species, and the universality of these effects has been found at both the molecular and phenotypic levels (Birchler et al., 1990; Devlin et al., 1982, 1988; Sun et al., 2013a,b,c; Hou et al., 2018; Johnson et al., 2020; Shi et al., 2021; Yang et al., 2021; Zhang S et al., 2021a).

Recent and the current results indicate a more intricate array of dosage effects beyond simply proportional effects of the varied genes. Genes located on the varied chromosomes may show a proportional gene dosage effect or compensation; the *trans*-acting effects of genes on unvaried chromosomes are more complicated and may present as negatively (inverse effect) or positively (direct effect) correlated with changed chromosomal regions, as well as an increase or decrease in both monosomy and trisomy (increased/decreased effect) (Guo and Birchler, 1994; Shi et al., 2021). It seems not surprising that changes of a chromosome or chromosomal fragments can result in global gene expression changes due to a disturbance through the gene regulatory networks and the dosage sensitivity of gene expression regulators. In *Drosophila* and maize any one gene product may be *trans* affected by multiple regions in the genome, and any one region may have a *trans* effect on many target genes (Birchler and Newton, 1981; Guo and Birchler, 1994; Birchler et al., 2001), which was also found in the present study.

Some studies have recognized the existence of dosage compensation or global aneuploid effects in human aneuploidy. Analysis of chromosome 21 microarray studies of Down syndrome revealed that approximately half of the transcripts were compensated (Ait Yahya-Graison et al., 2007). Another microarray study also found compensation in human autosomal trisomy cells (Altug-Teber et al., 2007). Whole-genome gene expression dysregulation domains have been observed in the study of trisomy 21 expression profiles (Letourneau et al., 2014). Among several sex chromosome aneuploidies, some X chromosome gene clusters were found to show nonlinear changes, and there was a close correlation between the XCI cluster and autosomal *trans*-regulation (Raznahan et al., 2018). Another study proposed that X chromosome dosage changes extend to the autosomes through the regulation of expression networks, which may share a common molecular mechanism (Zhang X et al., 2020). A recent study involving lncRNA and mRNA found that the expression of about one-third of X-linked genes on the active X chromosome (Xa) are *trans* affected by the number of inactive X chromosomes (Xi), and for almost all X-linked genes, the change in expression per Xi is less than that of Xa (San Roman et al., 2023). That study also analyzed the expression of genes on chromosome 21 in trisomy 21, and the median ΔE is 0.74, indicating that the gene expression on chromosome 21 is upregulated but not achieving full proportionality due to a compensation mechanism (San Roman et al., 2023).

Our study confirms that *trans*-acting effects across the genome extend to human. By generating ratio distributions and DEG analysis for *cis* and *trans* genes in human trisomies 13, 18, and 21, we found both dosage effects of 1.5-fold and dosage compensation for different sets of genes on the varied chromosomes. In the remainder of the genome both direct effects in proportion to the chromosome number changes and the inverse dosage effect were observed. We also found that the more imbalanced genotypes lead to stronger *trans*-acting effects, and a greater subset of *cis* genes showing compensation. Therefore, there seems to be a related mechanistic basis between the inverse dosage effect and dosage compensation as found in other organisms (Birchler, 1981; Birchler et al., 1990; Birchler and Veitia, 2021; Shi et al., 2021; Yang et al., 2021). In addition, we also analyzed the response of non-coding RNAs in human aneuploid cells and found that the expression of lncRNA is also affected. Previous studies of aneuploidy have mostly concentrated on mRNAs, but the inclusion here of lncRNAs in the study showed a strong effect.

A recent study concluded that most *cis* genes in human trisomies show a proportional gene dosage effect (Hwang et al., 2021). However, by analyzing the distributions of ratios of aneuploid/diploid levels for all expressed genes, it is obvious there are multiple peaks with some genes indeed exhibiting a proportional dosage effect but also a substantial subset being compensated and a still further subset reduced below the diploid with gradations between and beyond these peaks, with supporting evidence from DEG analysis. The same study claimed that members of multisubunit complexes are post transcriptionally compensated by protein degradation processes. While unincorporated subunits will indeed be degraded, this claim is inconsistent with the retention of dosage sensitive regulatory genes following whole genome duplications and their underrepresentation as duplicates in populations (Mottes et al., 2021; Zhang D et al., 2021), as well as their retention between heteromorphic sex chromosomes in three different vertebrate sex chromosome evolutions (Bellott et al., 2014, 2017; Bellott and Page, 2021). Further, it is inconsistent with disease states associated with dosage sensitive regulators (Ionita-Laza et al., 2009; Makino and McLysaght, 2010) given that transcription factors are generally dosage sensitive (Veitia, 2002; Seidman and Seidman, 2002). Our results showing global effects across the genome indicate they are effective in a dosage sensitive manner within cells.

There are different effects for sex chromosome and autosomal aneuploidies. We found that the addition or reduction of an X chromosome has little effect on the overall expression of the human genome, which likely involves the X inactivation mechanism and the fact that many dosage sensitive regulatory genes are retained between the X and Y chromosomes (Bellott et al., 2014). Sex chromosome divergence leads to the degeneration of one chromosome in the heterogametic sex to form a dosage scenario reminiscent of aneuploidy (Pessia et al., 2012). In order to maintain the balance between the single active sex chromosome and the diploid autosomes, as well as the balance between the expression of different numbers of sex chromosomes between males and females, a dosage compensation mechanism has evolved (Heard and Disteche, 2006). Therefore, in sex chromosome aneuploidy, the expression of X-linked genes is not greatly affected. The X inactivation mechanism not only accommodates a copy number change of dosage sensitive regulatory genes that might differ between the sex chromosomes (Pessia et al., 2012), but this also drastically reduces the *trans*-acting effects across the genome. With a substantial subset of genes in the autosomal trisomies showing compensation, this is related to the majority inverse effect in *trans* that also affects the varied chromosome. Yet, the reductions across the genome likely contribute to the detrimental effects of trisomies. The strong difference between autosomal and sex chromosomal effects illustrates the amelioration of these effects is a contributor to the evolution of sex chromosome dosage compensation.

The lesson from genome fractionation following whole genome duplications and the complementary pattern of gene classes in small scale duplications in populations, as noted above, is that the dosage of most types of genes is not particularly critical with the exception of those involved in multicomponent interactions, including those mediating gene expression. Thus, the driving force for the evolution of sex chromosome dosage compensation is to minimize the impact of those dosage sensitive genes not only on the sex chromosomes themselves (Pessia et al., 2012) but also for their impact across the whole genome. In summary, this study demonstrates that the *trans*-acting effects and compensation mechanisms are at work in human aneuploidy, and contribute to our understanding of gene expression regulation in unbalanced genomes and disease states.

## Materials and methods

### Key resources table

| Resource | Designation | Source or reference |
| --- | --- | --- |
| software, algorithm | R | http://www.R-project.org |
| software, algorithm | trim-galore v0.6.7 | https://www.bioinformatics.babraham.ac.uk/projects/trim_galore/ |
| software, algorithm | HISAT2 v2.2.1 | Kim et al., 2015 |
| software, algorithm | SAMtools v1.12 | Li et al., 2009 |
| software, algorithm | StringTie v2.1.5 | Pertea et al., 2015 |
| software, algorithm | prepDE.py | http://ccb.jhu.edu/software/stringtie/dl/prepDE.py3 |
| software, algorithm | ggplot2 v3.3.3 | Villanueva and Chen, 2019 |
| software, algorithm | clusterProfiler v4.0.2 | Wu et al., 2021 |
| software, algorithm | DESeq2 v1.32.0 | Love et al., 2014 |
| database | hTFtarget | Zhang Q et al., 2020 |
| database | CORUM | Giurgiu et al., 2019 |
| database | org.Hs.eg.db v3.13.0 | https://bioconductor.org/packages/org.Hs.eg.db/ |

### Data sets

The datasets of human aneuploidy analyzed in this study are from the Gene Expression Omnibus (GEO) public database, including: RNA-seq data of primary human fibroblasts in trisomies 13, 18, and 21 (GSE154418) (Hwang et al., 2021); RNA-seq data of primary peripheral blood mononuclear cells (PBMCs) derived from individuals with Turner syndrome (XO) and Klinefelter syndrome (XXY) (GSE126712) (Zhang X et al., 2020); RNA-seq data of primary skin fibroblasts from a pair of monozygotic twins discordant for trisomy 21, and from trisomy 21 and normal unrelated individuals (GSE55504) (Letourneau et al., 2014); RNA-seq data of fibroblasts from trisomy 21 individuals of different ages and sexes (GSE79842) (Sullivan et al., 2016); RNA-seq data of induced pluripotent stem cells (iPSCs) derived from primary fibroblast cell lines carrying XXY, XXXY, XXXXY, and normal sex chromosomes (GSE152001) (Astro et al., 2022).

### RNA sequencing raw data processing

Raw sequencing data were downloaded from the Sequence Read Archive (SRA) database (GSE154418, GSE126712) and analyzed using the HISAT2-StringTie pipeline (Pertea et al., 2016). The data were trimmed and filtered using trim-galore (version 0.6.7) (https://www.bioinformatics.babraham.ac.uk/projects/trim_galore/) to remove adaptors and low-quality sequences. RNA-seq reads were mapped to the human genome from Ensembl database (GRCh38 release 104) with the corresponding annotation using HISAT2 (version 2.2.1) (Kim et al., 2015) with the default parameters. Subsequently, SAMtools (version 1.12) (Li et al., 2009) was used to convert the alignment file format. Transcripts were assembled and quantitated using StringTie (version 2.1.5) (Pertea et al., 2015), and read counts were extracted using the Python script prepDE.py (http://ccb.jhu.edu/software/stringtie/dl/prepDE.py3). Three additional datasets of human aneuploidy were processed data obtained from the Gene Expression Omnibus (GEO) database (GSE55504, GSE79842, GSE152001).

### Ratio distribution

Read counts were normalized by dividing the total number of mapped reads in one sample to calculate CPM (Counts Per Million). The transcripts were filtered according to mean counts > 5 to avoid possible artifacts from low expression genes. Then the normalized data were averaged across genotypes, followed by calculating the ratio of each transcript by dividing the mean of aneuploid samples by the control samples. Ratio distribution plots were plotted in bins of 0.1 using ggplot2 package (version 3.3.3) (Villanueva and Chen, 2019) in the R program (R Core Team, 2022). The gene types belonging to “retained_intron”, “lincRNA”, “antisense”, “sense_intronic”, “sense_overlapping”, “3prime_overlapping_ncrna”, “bidirectional_promoter_lncrna”, “macro_lncRNA” in the annotation were defined as lncRNAs. The relationship between transcription factors and their targets was obtained from the hTFtarget database (Zhang Q et al., 2020). The genes of human macromolecular complexes were obtained from the core set in the CORUM database (Giurgiu et al., 2019).

### RNA sequencing analyses

Gene Ontology (GO) enrichment analysis was performed using the R package clusterProfiler (version 4.0.2) (Wu et al., 2021). The GO annotations were provided by org.Hs.eg.db (version 3.13.0), and GO functions in biological process were analyzed. Differential gene expression analysis was performed using DESeq2 (version 1.32.0) (Love et al., 2014). The transcrpts with mean counts < 10 were filtered out, and the threshold for differentially expressed genes was set as adjusted p-value < 0.05.

## Acknowledgements

Research was supported by National Natural Science Foundation of China (Grant No. 31871243 and No. 32070566) to L.S. and by USA National Science Foundation grant (IOS-1545780) to JB. We thank Reiner Veitia and Winston Bellott for comments on the manuscript.

## Additional information

## Funding

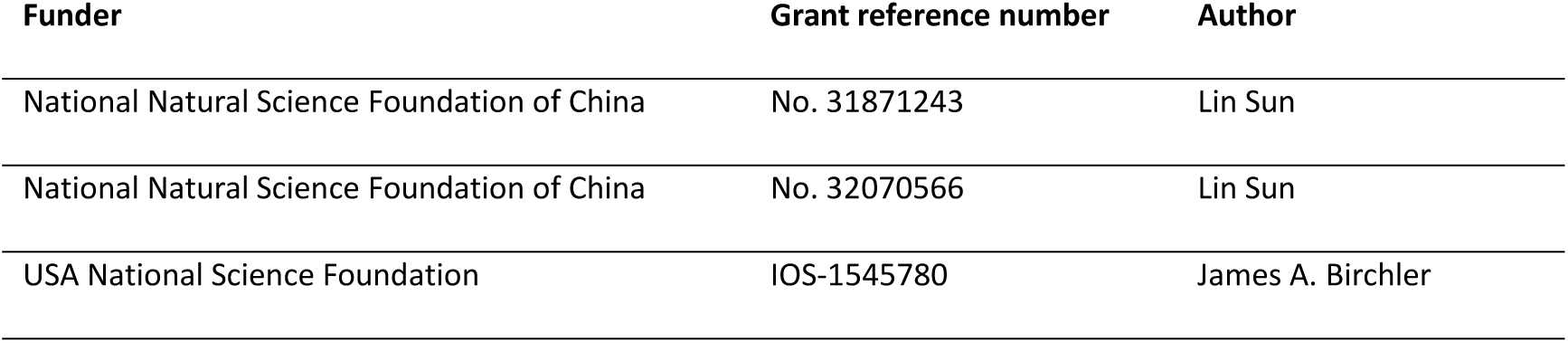

### Author contributions

Shuai Zhang: Methodology, Investigation, Data Curation, Formal analysis, Writing - Original Draft, Writing - Review & Editing, Visualization. Ruixue Wang: Methodology, Investigation, Formal analysis. Ludan Zhang: Methodology, Investigation, Validation,. James A. Birchler: Conceptualization, Writing - Original Draft, Writing - Review & Editing, Project administration, Funding acquisition. Lin Sun: Conceptualization, Writing - Original Draft, Writing - Review & Editing, Supervision, Project administration, Funding acquisition.

## Additional files

### Supplementary files

Supplementary File 1

### Data availability

The following previously published datasets were used:

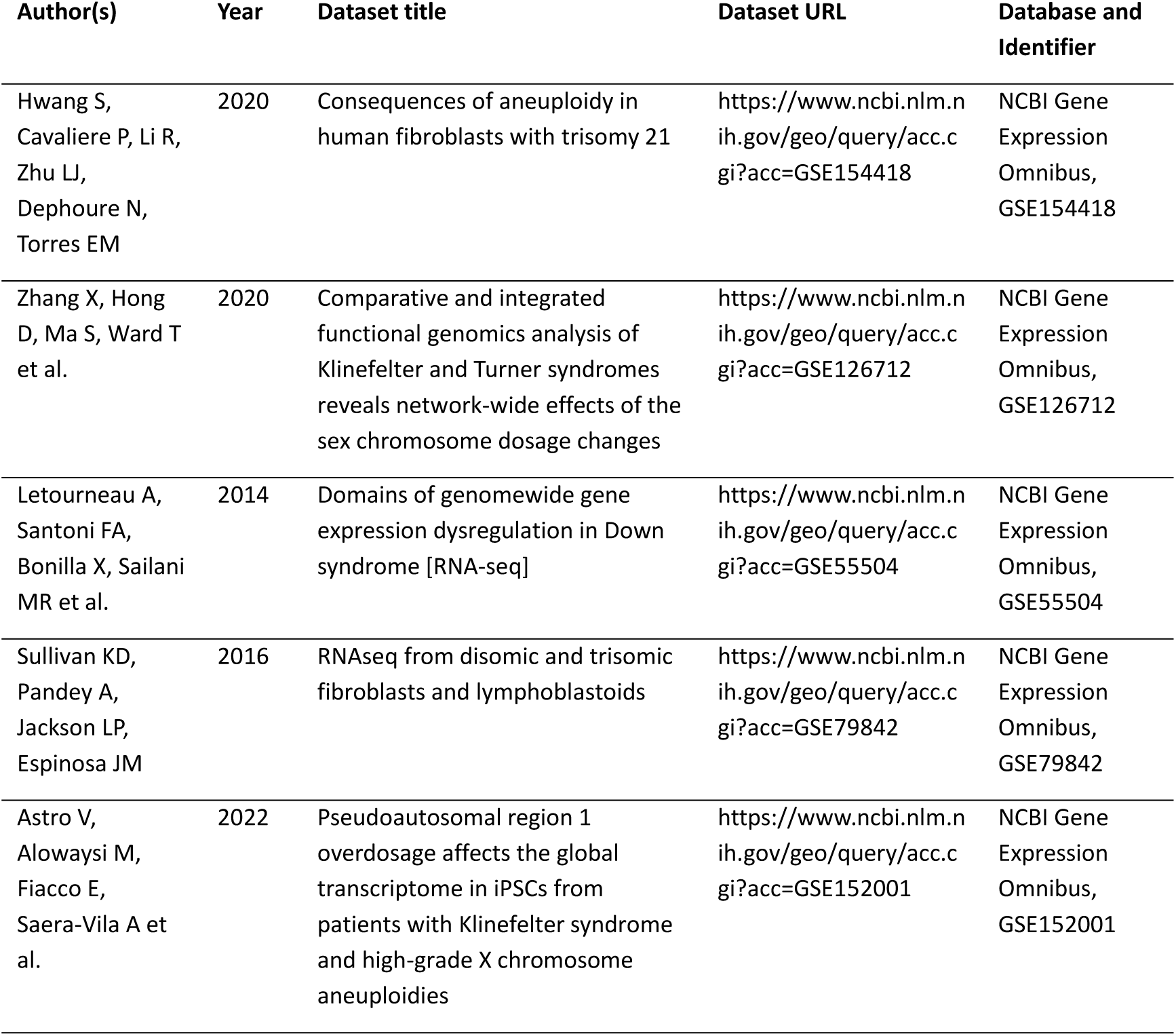

