## Supplementary File for "Inverse and Proportional *trans* Modulation of Gene Expression in Human Aneuploidies"

**Appendix 1—table 1.** The number of all differentially expressed genes (DEGs, adjusted p-value < 0.05) in human autosomal aneuploidy trisomy 13, trisomy 18, and trisomy 21.

| | | **all-transcripts** | | | **cis-transcripts** | | | **trans-transcripts** | | |
|---|---|---|---|---|---|---|---|---|---|---|
| | | **all** | **mRNA** | **lncRNA** | **all** | **mRNA** | **lncRNA** | **all** | **mRNA** | **lncRNA** |
| trisomy13 | up | 380 | 244 | 69 | 51 | 44 | 4 | 329 | 200 | 65 |
| | down | 1677 | 1304 | 120 | 14 | 12 | 1 | 1663 | 1292 | 119 |
| | normal | 45694 | 28191 | 10310 | 675 | 426 | 120 | 45019 | 27765 | 10190 |
| trisomy18 | up | 689 | 547 | 53 | 58 | 48 | 5 | 631 | 499 | 48 |
| | down | 1928 | 1340 | 285 | 10 | 9 | 0 | 1918 | 1331 | 285 |
| | normal | 45134 | 27852 | 10161 | 588 | 369 | 150 | 44546 | 27483 | 10011 |
| trisomy21 | up | 446 | 321 | 53 | 53 | 38 | 13 | 393 | 283 | 40 |
| | down | 320 | 233 | 31 | 0 | 0 | 0 | 320 | 233 | 31 |
| | normal | 46985 | 29185 | 10415 | 515 | 272 | 153 | 46470 | 28913 | 10262 |

**Appendix 1—table 2.** The number of low expression DEGs (mean counts > 5 and < 20, adjusted p-value < 0.05) in human autosomal aneuploidy trisomy 13, trisomy 18, and trisomy 21.

| | | **all-transcripts** | | | **cis-transcripts** | | | **trans-transcripts** | | |
|---|---|---|---|---|---|---|---|---|---|---|
| | | **all** | **mRNA** | **lncRNA** | **all** | **mRNA** | **lncRNA** | **all** | **mRNA** | **lncRNA** |
| trisomy13 | up | 38 | 14 | 16 | 1 | 1 | 0 | 37 | 13 | 16 |
| | down | 192 | 123 | 27 | 0 | 0 | 0 | 192 | 123 | 27 |
| | normal | 9341 | 3717 | 3311 | 137 | 57 | 27 | 9204 | 3660 | 3284 |
| trisomy18 | up | 43 | 22 | 11 | 2 | 1 | 1 | 41 | 21 | 10 |
| | down | 243 | 123 | 65 | 2 | 2 | 0 | 241 | 121 | 65 |
| | normal | 9285 | 3709 | 3278 | 118 | 51 | 42 | 9167 | 3658 | 3236 |
| trisomy21 | up | 63 | 38 | 13 | 2 | 0 | 2 | 61 | 38 | 11 |
| | down | 60 | 43 | 4 | 0 | 0 | 0 | 60 | 43 | 4 |
| | normal | 9448 | 3773 | 3337 | 109 | 39 | 47 | 9339 | 3734 | 3290 |

**Appendix 1—table 3.** The number of medium expression DEGs (mean counts > 20 and < 100, adjusted p-value < 0.05) in human autosomal aneuploidy trisomy 13, trisomy 18, and trisomy 21.

| | | **all-transcripts** | | | **cis-transcripts** | | | **trans-transcripts** | | |
|---|---|---|---|---|---|---|---|---|---|---|
| | | **all** | **mRNA** | **lncRNA** | **all** | **mRNA** | **lncRNA** | **all** | **mRNA** | **lncRNA** |
| trisomy13 | up | 200 | 129 | 37 | 9 | 7 | 2 | 191 | 122 | 35 |

| | | | | | | | | | | |
|---|---|---|---|---|---|---|---|---|---|---|
| | down | 725 | 534 | 55 | 9 | 8 | 1 | 716 | 526 | 54 |
| | normal | 15816 | 7822 | 4818 | 233 | 119 | 65 | 15583 | 7703 | 4753 |
| trisomy18 | up | 232 | 161 | 25 | 9 | 5 | 3 | 223 | 156 | 22 |
| | down | 812 | 512 | 132 | 5 | 4 | 0 | 807 | 508 | 132 |
| | normal | 15697 | 7812 | 4753 | 216 | 111 | 79 | 15481 | 7701 | 4674 |
| trisomy21 | up | 221 | 153 | 24 | 12 | 4 | 7 | 209 | 149 | 17 |
| | down | 146 | 101 | 15 | 0 | 0 | 0 | 146 | 101 | 15 |
| | normal | 16374 | 8231 | 4871 | 199 | 83 | 72 | 16175 | 8148 | 4799 |

**Appendix 1—table 4.** The number of high expression DEGs (mean counts > 100, adjusted p-value < 0.05) in human autosomal aneuploidy trisomy 13, trisomy 18, and trisomy 21.

| | | **all-transcripts** | | | **cis-transcripts** | | | **trans-transcripts** | | |
|---|---|---|---|---|---|---|---|---|---|---|
| | | **all** | **mRNA** | **lncRNA** | **all** | **mRNA** | **lncRNA** | **all** | **mRNA** | **lncRNA** |
| trisomy13 | up | 142 | 101 | 16 | 41 | 36 | 2 | 101 | 65 | 14 |
| | down | 760 | 647 | 38 | 5 | 4 | 0 | 755 | 643 | 38 |
| | normal | 20537 | 16652 | 2181 | 305 | 250 | 28 | 20232 | 16402 | 2153 |
| trisomy18 | up | 414 | 364 | 17 | 47 | 42 | 1 | 367 | 322 | 16 |
| | down | 873 | 705 | 88 | 3 | 3 | 0 | 870 | 702 | 88 |
| | normal | 20152 | 16331 | 2130 | 254 | 207 | 29 | 19898 | 16124 | 2101 |
| trisomy21 | up | 162 | 130 | 16 | 39 | 34 | 4 | 123 | 96 | 12 |
| | down | 114 | 89 | 12 | 0 | 0 | 0 | 114 | 89 | 12 |
| | normal | 21163 | 17181 | 2207 | 207 | 150 | 34 | 20956 | 17031 | 2173 |

**Appendix 1—table 5.** The number of significantly up-regulated and down-regulated transcripts (adjusted p-value < 0.05) in human sex chromosome aneuploidy XO and XXY.

| | | **all-transcripts** | | | **X-linked transcripts** | | | **autosomal transcripts** | | |
|---|---|---|---|---|---|---|---|---|---|---|
| | | **all** | **mRNA** | **lncRNA** | **all** | **mRNA** | **lncRNA** | **all** | **mRNA** | **lncRNA** |
| XO/XX | up | 280 | 201 | 46 | 12 | 11 | 1 | 268 | 190 | 45 |
| | down | 330 | 220 | 43 | 32 | 16 | 9 | 298 | 204 | 34 |
| | normal | 43347 | 25078 | 10927 | 1224 | 823 | 173 | 42088 | 24239 | 10747 |
| XXY/XY | up | 119 | 79 | 14 | 24 | 10 | 8 | 94 | 69 | 6 |
| | down | 102 | 78 | 8 | 3 | 2 | 0 | 99 | 76 | 8 |
| | normal | 43736 | 25342 | 10994 | 1241 | 838 | 175 | 42461 | 24488 | 10812 |
| XXY/XX | up | 129 | 87 | 19 | 3 | 3 | 0 | 94 | 69 | 12 |
| | down | 133 | 92 | 12 | 3 | 1 | 0 | 130 | 91 | 12 |

| | | | | | | | | | | |
|---|---|---|---|---|---|---|---|---|---|---|
| | normal | 43695 | 25320 | 10985 | 1262 | 846 | 183 | 42430 | 24473 | 10802 |
| XY/XX | up | 130 | 90 | 17 | 0 | 0 | 0 | 99 | 76 | 10 |
| | down | 179 | 120 | 28 | 19 | 7 | 8 | 160 | 113 | 20 |
| | normal | 43648 | 25289 | 10971 | 1249 | 843 | 175 | 42395 | 24444 | 10796 |

**Appendix 1—table 6.** The proportion of up-regulated (ratio > 1.25) and down-regulated (ratio < 0.8) transcripts on X chromosome and autosomes in human sex chromosome aneuploidy XXY, XXXY, and XXXXY.

| | | **down-regulated (ratio < 0.8)** | | | | **up-regulated (ratio > 1.25)** | | | |
|---|---|---|---|---|---|---|---|---|---|
| | | XX/XY | XXY/XY | XXXY/XY | XXXXY/XY | XX/XY | XXY/XY | XXXY/XY | XXXXY/XY |
| all type | chrX | 23.60% | 13.76% | 37.27% | 29.21% | 28.46% | 24.44% | 22.38% | 29.78% |
| | autosomes | 24.85% | 13.96% | 33.68% | 28.67% | 24.41% | 22.13% | 25.64% | 25.93% |
| mRNA | chrX | 15.21% | 9.87% | 26.70% | 25.89% | 22.98% | 11.97% | 19.09% | 21.84% |
| | autosomes | 15.69% | 8.54% | 23.18% | 21.51% | 18.40% | 10.39% | 21.78% | 18.72% |
| lncRNA | chrX | 24.91% | 13.75% | 38.29% | 31.97% | 21.19% | 20.45% | 21.93% | 23.05% |
| | autosomes | 22.36% | 11.58% | 29.90% | 27.08% | 22.55% | 20.04% | 26.11% | 23.63% |

**Appendix 1—table 7.** The number of significantly up-regulated and down-regulated transcripts (t-test p-value < 0.05) in human sex chromosome aneuploidy XXY, XXXY, and XXXXY.

| | | **all-transcripts** | | | **X-linked transcripts** | | | **autosomal transcripts** | | |
|---|---|---|---|---|---|---|---|---|---|---|
| | | **all** | **mRNA** | **lncRNA** | **all** | **mRNA** | **lncRNA** | **all** | **mRNA** | **lncRNA** |
| XX/XY | up | 3160 | 2475 | 1697 | 136 | 109 | 38 | 2999 | 2364 | 1659 |
| | down | 2974 | 1915 | 1580 | 122 | 93 | 32 | 2778 | 1807 | 1539 |
| | normal | 23114 | 11699 | 11720 | 822 | 416 | 201 | 21907 | 11261 | 11519 |
| XXY/XY | up | 4725 | 2818 | 2589 | 184 | 110 | 44 | 4476 | 2705 | 2543 |
| | down | 2118 | 1785 | 1142 | 117 | 112 | 33 | 1987 | 1672 | 1109 |
| | normal | 22694 | 11488 | 11311 | 788 | 396 | 195 | 21487 | 11057 | 11109 |
| XXXY/XY | up | 2062 | 1326 | 1148 | 59 | 45 | 14 | 1980 | 1280 | 1134 |
| | down | 4122 | 2110 | 2043 | 167 | 106 | 46 | 3885 | 2000 | 1996 |
| | normal | 22981 | 12652 | 11806 | 848 | 467 | 211 | 21748 | 12151 | 11587 |
| XXXXY/XY | up | 3534 | 2816 | 1958 | 143 | 107 | 36 | 3371 | 2707 | 1922 |
| | down | 4401 | 3146 | 2481 | 198 | 171 | 63 | 4143 | 2973 | 2418 |
| | normal | 21319 | 10127 | 10567 | 738 | 340 | 172 | 20179 | 9752 | 10386 |

**Appendix 1—table 8.** The number of up- and down-regulated (padj < 0.1) target genes of all transcription factors (TFs) in human autosomal aneuploidy trisomy 13, trisomy 18, and trisomy 21.

| | | **cis-TF (all)** | | **trans-TF (all)** | |
|---|---|---|---|---|---|
| | | **cis-target** | **trans-target** | **cis-target** | **trans-target** |
| trisomy 13 | up | 44 | 344 | 61 | 426 |
| | down | 12 | 1432 | 17 | 1964 |
| | normal | 346 | 30721 | 583 | 40278 |
| trisomy 18 | up | 2 | 36 | 70 | 830 |
| | down | 0 | 119 | 11 | 2277 |
| | normal | 28 | 2181 | 529 | 40328 |
| trisomy 21 | up | 53 | 394 | 53 | 440 |
| | down | 0 | 332 | 0 | 375 |
| | normal | 377 | 38837 | 419 | 42720 |

**Appendix 1—table 9.** The number of up- and down-regulated (padj < 0.1) target genes of positively changed TFs (> 1.25) in human autosomal aneuploidy trisomy 13, trisomy 18, and trisomy 21.

| | | **cis-TF (> 1.25)** | | **trans-TF (> 1.25)** | |
|---|---|---|---|---|---|
| | | **cis-target** | **trans-target** | **cis-target** | **trans-target** |
| trisomy 13 | up | 42 | 333 | 60 | 425 |
| | down | 12 | 1373 | 17 | 1938 |
| | normal | 334 | 29765 | 576 | 39938 |
| trisomy 18 | up | 2 | 33 | 70 | 826 |
| | down | 0 | 110 | 11 | 2264 |
| | normal | 28 | 2078 | 526 | 40165 |
| trisomy 21 | up | 53 | 394 | 51 | 434 |
| | down | 0 | 332 | 0 | 366 |
| | normal | 377 | 38837 | 406 | 42247 |

**Appendix 1—table 10.** The number of up- and down-regulated (padj < 0.1) target genes of negatively changed TFs (< 0.8) in human autosomal aneuploidy trisomy 13, trisomy 18, and

trisomy 21.

| | | cis-TF (< 0.8) | | trans-TF (< 0.8) | |
|---|---|---|---|---|---|
| | | **cis-target** | **trans-target** | **cis-target** | **trans-target** |
| trisomy 13 | up | 10 | 165 | 61 | 424 |
| | down | 4 | 620 | 17 | 1948 |
| | normal | 119 | 13981 | 580 | 40110 |
| trisomy 18 | up | 2 | 31 | 70 | 827 |
| | down | 0 | 105 | 11 | 2264 |
| | normal | 28 | 1903 | 529 | 40163 |
| trisomy 21 | up | 12 | 103 | 52 | 434 |
| | down | 0 | 91 | 0 | 368 |
| | normal | 99 | 9836 | 417 | 42446 |

**Appendix 1—table 11.** The number of up- and down-regulated (padj < 0.1) target genes of unchanged TFs (> 0.8 and < 1.25) in human autosomal aneuploidy trisomy 13, trisomy 18, and trisomy 21.

| | | cis-TF (>0.8 & <1.25) | | trans-TF (>0.8 & <1.25) | |
|---|---|---|---|---|---|
| | | **cis-target** | **trans-target** | **cis-target** | **trans-target** |
| trisomy 13 | up | 1 | 42 | 61 | 425 |
| | down | 1 | 148 | 17 | 1948 |
| | normal | 30 | 3371 | 583 | 40037 |
| trisomy 18 | up | 2 | 27 | 70 | 820 |
| | down | 0 | 94 | 11 | 2254 |
| | normal | 28 | 1659 | 525 | 40028 |
| trisomy 21 | up | 42 | 355 | 52 | 437 |
| | down | 0 | 296 | 0 | 374 |
| | normal | 329 | 35032 | 417 | 42572 |

**Appendix 1—table 12.** The results of statistical tests.

| figures | distribution or comparison | test | p-value |
|---|---|---|---|
| Fig. 1A | trisomy13 cis | normality test | 0.0000E+00 |
| Fig. 1B | trisomy13 trans | normality test | 0.0000E+00 |
| Fig. 1C | trisomy18 cis | normality test | 0.0000E+00 |
| Fig. 1D | trisomy18 trans | normality test | 0.0000E+00 |

| | | | |
|---|---|---|---|
| Fig. 1E | trisomy21 cis | normality test | 0.0000E+00 |
| Fig. 1F | trisomy21 trans | normality test | 0.0000E+00 |
| Fig. 1-fig. S6A | trisomy13 cis - low vs high | K-S test | 1.9635E-02 |
| Fig. 1-fig. S6B | trisomy13 trans - low vs high | K-S test | 0.0000E+00 |
| Fig. 1-fig. S6C | trisomy18 cis - low vs high | K-S test | 1.1859E-04 |
| Fig. 1-fig. S6D | trisomy18 trans - low vs high | K-S test | 0.0000E+00 |
| Fig. 1-fig. S6E | trisomy21 cis - low vs high | K-S test | 3.1638E-02 |
| Fig. 1-fig. S6F | trisomy21 trans - low vs high | K-S test | 0.0000E+00 |
| Fig. 1-fig. S6A | trisomy13 cis - medium vs high | K-S test | 7.7676E-02 |
| Fig. 1-fig. S6B | trisomy13 trans - medium vs high | K-S test | 0.0000E+00 |
| Fig. 1-fig. S6C | trisomy18 cis - medium vs high | K-S test | 4.7205E-02 |
| Fig. 1-fig. S6D | trisomy18 trans - medium vs high | K-S test | 0.0000E+00 |
| Fig. 1-fig. S6E | trisomy21 cis - medium vs high | K-S test | 8.9044E-02 |
| Fig. 1-fig. S6F | trisomy21 trans - medium vs high | K-S test | 0.0000E+00 |
| Fig. 1-fig. S6A | trisomy13 cis - low vs medium | K-S test | 1.9635E-02 |
| Fig. 1-fig. S6B | trisomy13 trans - low vs medium | K-S test | 0.0000E+00 |
| Fig. 1-fig. S6C | trisomy18 cis - low vs medium | K-S test | 1.1859E-04 |
| Fig. 1-fig. S6D | trisomy18 trans - low vs medium | K-S test | 0.0000E+00 |
| Fig. 1-fig. S6E | trisomy21 cis - low vs medium | K-S test | 3.1638E-02 |
| Fig. 1-fig. S6F | trisomy21 trans - low vs medium | K-S test | 0.0000E+00 |
| Fig. 2A | trisomy18/diploid - DC genes identified in trisomy13 vs all chr13 genes | K-S test | 4.9869E-01 |
| Fig. 2B | trisomy21/diploid - DC genes identified in trisomy13 vs all chr13 genes | K-S test | 4.6560E-01 |
| Fig. 2C | trisomy13/diploid - DC genes identified in trisomy18 vs all chr18 genes | K-S test | 5.5409E-02 |
| Fig. 2D | trisomy21/diploid - DC genes identified in trisomy18 vs all chr18 genes | K-S test | 1.2115E-02 |
| Fig. 2E | trisomy13/diploid - DC genes identified in trisomy21 vs all chr21 genes | K-S test | 2.3720E-03 |
| Fig. 2F | trisomy18/diploid - DC genes identified in trisomy21 vs all chr21 genes | K-S test | 4.1489E-01 |
| Fig. 2J | trisomy13/diploid - IDE genes identified in trisomy 18 and 21 vs all chr13 genes | K-S test | 2.9480E-03 |
| Fig. 2K | trisomy18/diploid - IDE genes identified in trisomy 13 and 21 vs all chr18 genes | K-S test | 2.3487E-06 |
| Fig. 2L | trisomy21/diploid - IDE genes identified in trisomy 13 and 18 vs all chr21 genes | K-S test | 1.8613E-03 |
| Fig. 2A | trisomy18/diploid - DC genes identified in trisomy13 | normality test | 1.3650E-05 |
| Fig. 2B | trisomy21/diploid - DC genes identified in trisomy13 | normality test | 1.1770E-01 |
| Fig. 2C | trisomy13/diploid - DC genes identified in trisomy18 | normality test | 5.3040E-07 |
| Fig. 2D | trisomy21/diploid - DC genes identified in trisomy18 | normality test | 1.1370E-01 |
| Fig. 2E | trisomy13/diploid - DC genes identified in trisomy21 | normality test | 2.8470E-01 |

| | | | |
|---|---|---|---|
| Fig. 2F | trisomy18/diploid - DC genes identified in trisomy21 | normality test | 6.4590E-01 |
| Fig. 2J | trisomy13/diploid - IDE genes identified in trisomy 18 and 21 | normality test | 7.0380E-09 |
| Fig. 2K | trisomy18/diploid - IDE genes identified in trisomy 13 and 21 | normality test | 8.4840E-12 |
| Fig. 2L | trisomy21/diploid - IDE genes identified in trisomy 13 and 18 | normality test | 1.3410E-06 |
| Fig. 3A | trisomy21 twins cis | normality test | 6.1850E-08 |
| Fig. 3B | trisomy21 twins trans | normality test | 0.0000E+00 |
| Fig. 3C | trisomy21 unrelated individuals cis | normality test | 2.9980E-15 |
| Fig. 3D | trisomy21 unrelated individuals trans | normality test | 0.0000E+00 |
| Fig. 3E | trisomy21 female 2-3 days old cis | normality test | 0.0000E+00 |
| Fig. 3F | trisomy21 female 2-3 days old trans | normality test | 0.0000E+00 |
| Fig. 3G | trisomy21 male 1-2 years old cis | normality test | 0.0000E+00 |
| Fig. 3H | trisomy21 male 1-2 years old trans | normality test | 0.0000E+00 |
| Fig. 3I | trisomy21 female 11-19 years old cis | normality test | 2.6610E-11 |
| Fig. 3J | trisomy21 female 11-19 years old trans | normality test | 0.0000E+00 |
| Fig. 3K | trisomy21 male 19-21 years old cis | normality test | 9.9250E-14 |
| Fig. 3L | trisomy21 male 19-21 years old trans | normality test | 0.0000E+00 |
| Fig. 3A-C | trisomy21 cis - twins vs unrelated individuals | K-S test | 2.9615E-08 |
| Fig. 3B-D | trisomy21 trans - twins vs unrelated individuals | K-S test | 0.0000E+00 |
| Fig. 3E-G | trisomy21 cis - female 2-3 days vs male 1-2 years | K-S test | 5.5823E-02 |
| Fig. 3F-H | trisomy21 trans - female 2-3 days vs male 1-2 years | K-S test | 0.0000E+00 |
| Fig. 3E-I | trisomy21 cis - female 2-3 days vs female 11-19 years | K-S test | 2.0190E-05 |
| Fig. 3F-J | trisomy21 trans - female 2-3 days vs female 11-19 years | K-S test | 0.0000E+00 |
| Fig. 3E-K | trisomy21 cis - female 2-3 days vs male 19-21 years | K-S test | 4.1559E-05 |
| Fig. 3F-L | trisomy21 trans - female 2-3 days vs male 19-21 years | K-S test | 0.0000E+00 |
| Fig. 3G-I | trisomy21 cis - male 1-2 years vs female 11-19 years | K-S test | 2.7751E-03 |
| Fig. 3H-J | trisomy21 trans - male 1-2 years vs female 11-19 years | K-S test | 0.0000E+00 |
| Fig. 3G-K | trisomy21 cis - male 1-2 years vs male 19-21 years | K-S test | 1.7902E-03 |
| Fig. 3H-L | trisomy21 trans - male 1-2 years vs male 19-21 years | K-S test | 0.0000E+00 |
| Fig. 3I-K | trisomy21 cis - female 11-19 years vs male 19-21 years | K-S test | 8.7020E-01 |
| Fig. 3J-L | trisomy21 trans - female 11-19 years vs male 19-21 years | K-S test | 2.2204E-16 |
| Fig. 4A | trisomy13 cis - mRNA vs lncRNA | K-S test | 1.8172E-01 |
| Fig. 4B | trisomy13 trans - mRNA vs lncRNA | K-S test | 0.0000E+00 |
| Fig. 4C | trisomy18 cis - mRNA vs lncRNA | K-S test | 9.0784E-05 |
| Fig. 4D | trisomy18 trans - mRNA vs lncRNA | K-S test | 0.0000E+00 |
| Fig. 4E | trisomy21 cis - mRNA vs lncRNA | K-S test | 1.7831E-01 |
| Fig. 4F | trisomy21 trans - mRNA vs lncRNA | K-S test | 6.6613E-15 |
| Fig. 4-fig. S7A | trisomy13 cis mRNA - low vs high | K-S test | 6.3714E-03 |
| Fig. 4-fig. S7C | trisomy18 cis mRNA - low vs high | K-S test | 2.5040E-04 |

| | | | |
|---|---|---|---|
| Fig. 4-fig. S7E | trisomy21 cis mRNA - low vs high | K-S test | 8.0375E-04 |
| Fig. 4-fig. S7A | trisomy13 cis mRNA - medium vs high | K-S test | 1.1507E-07 |
| Fig. 4-fig. S7C | trisomy18 cis mRNA - medium vs high | K-S test | 1.8874E-15 |
| Fig. 4-fig. S7E | trisomy21 cis mRNA - medium vs high | K-S test | 1.4906E-08 |
| Fig. 4-fig. S7A | trisomy13 cis mRNA - low vs medium | K-S test | 7.7870E-03 |
| Fig. 4-fig. S7C | trisomy18 cis mRNA - low vs medium | K-S test | 4.2391E-03 |
| Fig. 4-fig. S7E | trisomy21 cis mRNA - low vs medium | K-S test | 1.8537E-02 |
| Fig. 4-fig. S7B | trisomy13 cis lncRNA - low vs high | K-S test | 5.4989E-02 |
| Fig. 4-fig. S7D | trisomy18 cis lncRNA - low vs high | K-S test | 4.2252E-02 |
| Fig. 4-fig. S7F | trisomy21 cis lncRNA - low vs high | K-S test | 5.1557E-01 |
| Fig. 4-fig. S7B | trisomy13 cis lncRNA - medium vs high | K-S test | 3.0545E-01 |
| Fig. 4-fig. S7D | trisomy18 cis lncRNA - medium vs high | K-S test | 4.4189E-02 |
| Fig. 4-fig. S7F | trisomy21 cis lncRNA - medium vs high | K-S test | 5.7932E-01 |
| Fig. 4-fig. S7B | trisomy13 cis lncRNA - low vs medium | K-S test | 4.0345E-06 |
| Fig. 4-fig. S7D | trisomy18 cis lncRNA - low vs medium | K-S test | 4.0950E-02 |
| Fig. 4-fig. S7F | trisomy21 cis lncRNA - low vs medium | K-S test | 4.5027E-07 |
| Fig. 4-fig. S7A-B | trisomy13 cis low - mRNA vs lncRNA | K-S test | 4.2754E-04 |
| Fig. 4-fig. S7C-D | trisomy18 cis low - mRNA vs lncRNA | K-S test | 1.2166E-03 |
| Fig. 4-fig. S7E-F | trisomy21 cis low - mRNA vs lncRNA | K-S test | 1.1140E-08 |
| Fig. 4-fig. S7A-B | trisomy13 cis medium - mRNA vs lncRNA | K-S test | 7.4920E-02 |
| Fig. 4-fig. S7C-D | trisomy18 cis medium - mRNA vs lncRNA | K-S test | 2.1461E-03 |
| Fig. 4-fig. S7E-F | trisomy21 cis medium - mRNA vs lncRNA | K-S test | 9.5764E-02 |
| Fig. 4-fig. S7A-B | trisomy13 cis high - mRNA vs lncRNA | K-S test | 4.4374E-03 |
| Fig. 4-fig. S7C-D | trisomy18 cis high - mRNA vs lncRNA | K-S test | 2.6918E-03 |
| Fig. 4-fig. S7E-F | trisomy21 cis high - mRNA vs lncRNA | K-S test | 9.7453E-04 |
| Fig. 4-fig. S8A | trisomy13 trans mRNA - low vs high | K-S test | 0.0000E+00 |
| Fig. 4-fig. S8C | trisomy18 trans mRNA - low vs high | K-S test | 0.0000E+00 |
| Fig. 4-fig. S8E | trisomy21 trans mRNA - low vs high | K-S test | 0.0000E+00 |
| Fig. 4-fig. S8A | trisomy13 trans mRNA - medium vs high | K-S test | 0.0000E+00 |
| Fig. 4-fig. S8C | trisomy18 trans mRNA - medium vs high | K-S test | 0.0000E+00 |
| Fig. 4-fig. S8E | trisomy21 trans mRNA - medium vs high | K-S test | 0.0000E+00 |
| Fig. 4-fig. S8A | trisomy13 trans mRNA - low vs medium | K-S test | 0.0000E+00 |
| Fig. 4-fig. S8C | trisomy18 trans mRNA - low vs medium | K-S test | 0.0000E+00 |
| Fig. 4-fig. S8E | trisomy21 trans mRNA - low vs medium | K-S test | 0.0000E+00 |
| Fig. 4-fig. S8B | trisomy13 trans lncRNA - low vs high | K-S test | 3.9857E-14 |
| Fig. 4-fig. S8D | trisomy18 trans lncRNA - low vs high | K-S test | 0.0000E+00 |
| Fig. 4-fig. S8F | trisomy21 trans lncRNA - low vs high | K-S test | 0.0000E+00 |
| Fig. 4-fig. S8B | trisomy13 trans lncRNA - medium vs high | K-S test | 5.9286E-14 |
| Fig. 4-fig. S8D | trisomy18 trans lncRNA - medium vs high | K-S test | 0.0000E+00 |
| Fig. 4-fig. S8F | trisomy21 trans lncRNA - medium vs high | K-S test | 0.0000E+00 |
| Fig. 4-fig. S8B | trisomy13 trans lncRNA - low vs medium | K-S test | 0.0000E+00 |
| Fig. 4-fig. S8D | trisomy18 trans lncRNA - low vs medium | K-S test | 0.0000E+00 |
| Fig. 4-fig. S8F | trisomy21 trans lncRNA - low vs medium | K-S test | 0.0000E+00 |

| | | | |
|---|---|---|---|
| Fig. 4-fig. S8A-B | trisomy13 trans low - mRNA vs lncRNA | K-S test | 0.0000E+00 |
| Fig. 4-fig. S8C-D | trisomy18 trans low - mRNA vs lncRNA | K-S test | 0.0000E+00 |
| Fig. 4-fig. S8E-F | trisomy21 trans low - mRNA vs lncRNA | K-S test | 0.0000E+00 |
| Fig. 4-fig. S8A-B | trisomy13 trans medium - mRNA vs lncRNA | K-S test | 0.0000E+00 |
| Fig. 4-fig. S8C-D | trisomy18 trans medium - mRNA vs lncRNA | K-S test | 0.0000E+00 |
| Fig. 4-fig. S8E-F | trisomy21 trans medium - mRNA vs lncRNA | K-S test | 0.0000E+00 |
| Fig. 4-fig. S8A-B | trisomy13 trans high - mRNA vs lncRNA | K-S test | 0.0000E+00 |
| Fig. 4-fig. S8C-D | trisomy18 trans high - mRNA vs lncRNA | K-S test | 0.0000E+00 |
| Fig. 4-fig. S8E-F | trisomy21 trans high - mRNA vs lncRNA | K-S test | 0.0000E+00 |
| Fig. 5A | chrX all genes - XO/XX vs XY/XX | K-S test | 3.5310E-03 |
| Fig. 5A | chrX all genes - XO/XX vs XXY/XX | K-S test | 8.9655E-06 |
| Fig. 5A | chrX all genes - XY/XX vs XXY/XX | K-S test | 7.8900E-06 |
| Fig. 5A | chrX all genes - XO/XX vs XXY/XY | K-S test | 9.1982E-07 |
| Fig. 5A | chrX all genes - XY/XX vs XXY/XY | K-S test | 1.7250E-10 |
| Fig. 5B | autosome all genes - XO/XX vs XY/XX | K-S test | 0.0000E+00 |
| Fig. 5B | autosome all genes - XO/XX vs XXY/XX | K-S test | 0.0000E+00 |
| Fig. 5B | autosome all genes - XY/XX vs XXY/XX | K-S test | 0.0000E+00 |
| Fig. 5B | autosome all genes - XO/XX vs XXY/XY | K-S test | 0.0000E+00 |
| Fig. 5B | autosome all genes - XY/XX vs XXY/XY | K-S test | 0.0000E+00 |
| Fig. 5C | chrX mRNA - XO/XX vs XY/XX | K-S test | 4.7309E-02 |
| Fig. 5C | chrX mRNA - XO/XX vs XXY/XX | K-S test | 2.4176E-01 |
| Fig. 5C | chrX mRNA - XY/XX vs XXY/XX | K-S test | 2.1037E-01 |
| Fig. 5C | chrX mRNA - XO/XX vs XXY/XY | K-S test | 3.7891E-01 |
| Fig. 5C | chrX mRNA - XY/XX vs XXY/XY | K-S test | 1.6563E-01 |
| Fig. 5D | autosome mRNA - XO/XX vs XY/XX | K-S test | 3.3307E-16 |
| Fig. 5D | autosome mRNA - XO/XX vs XXY/XX | K-S test | 9.5024E-13 |
| Fig. 5D | autosome mRNA - XY/XX vs XXY/XX | K-S test | 1.3433E-02 |
| Fig. 5D | autosome mRNA - XO/XX vs XXY/XY | K-S test | 0.0000E+00 |
| Fig. 5D | autosome mRNA - XY/XX vs XXY/XY | K-S test | 0.0000E+00 |
| Fig. 5E | chrX lncRNA - XO/XX vs XY/XX | K-S test | 2.9243E-01 |
| Fig. 5E | chrX lncRNA - XO/XX vs XXY/XX | K-S test | 2.0961E-05 |
| Fig. 5E | chrX lncRNA - XY/XX vs XXY/XX | K-S test | 1.5809E-05 |
| Fig. 5E | chrX lncRNA - XO/XX vs XXY/XY | K-S test | 1.5674E-08 |
| Fig. 5E | chrX lncRNA - XY/XX vs XXY/XY | K-S test | 5.8376E-13 |
| Fig. 5F | autosome lncRNA - XO/XX vs XY/XX | K-S test | 4.4020E-12 |
| Fig. 5F | autosome lncRNA - XO/XX vs XXY/XX | K-S test | 0.0000E+00 |
| Fig. 5F | autosome lncRNA - XY/XX vs XXY/XX | K-S test | 0.0000E+00 |
| Fig. 5F | autosome lncRNA - XO/XX vs XXY/XY | K-S test | 0.0000E+00 |
| Fig. 5F | autosome lncRNA - XY/XX vs XXY/XY | K-S test | 0.0000E+00 |
| Fig. 5G | chrX all genes - XX/XY vs XXY/XY | K-S test | 1.2821E-07 |
| Fig. 5G | chrX all genes - XXXY/XY vs XXY/XY | K-S test | 0.0000E+00 |
| Fig. 5G | chrX all genes - XXXXY/XY vs XXY/XY | K-S test | 6.6613E-16 |
| Fig. 5G | chrX all genes - XXXY/XY vs XXXXY/XY | K-S test | 4.4195E-05 |

| | | | |
|---|---|---|---|
| Fig. 5H | autosome all genes - XX/XY vs XXY/XY | K-S test | 0.0000E+00 |
| Fig. 5H | autosome all genes - XXXY/XY vs XXY/XY | K-S test | 0.0000E+00 |
| Fig. 5H | autosome all genes - XXXXY/XY vs XXY/XY | K-S test | 0.0000E+00 |
| Fig. 5H | autosome all genes - XXXY/XY vs XXXXY/XY | K-S test | 0.0000E+00 |
| Fig. 5I | chrX mRNA - XX/XY vs XXY/XY | K-S test | 1.7458E-04 |
| Fig. 5I | chrX mRNA - XXXY/XY vs XXY/XY | K-S test | 1.0285E-10 |
| Fig. 5I | chrX mRNA - XXXXY/XY vs XXY/XY | K-S test | 6.1279E-12 |
| Fig. 5I | chrX mRNA - XXXY/XY vs XXXXY/XY | K-S test | 3.7880E-01 |
| Fig. 5J | autosome mRNA - XX/XY vs XXY/XY | K-S test | 0.0000E+00 |
| Fig. 5J | autosome mRNA - XXXY/XY vs XXY/XY | K-S test | 0.0000E+00 |
| Fig. 5J | autosome mRNA - XXXXY/XY vs XXY/XY | K-S test | 0.0000E+00 |
| Fig. 5J | autosome mRNA - XXXY/XY vs XXXXY/XY | K-S test | 1.4113E-07 |
| Fig. 5K | chrX lncRNA - XX/XY vs XXY/XY | K-S test | 2.8385E-03 |
| Fig. 5K | chrX lncRNA - XXXY/XY vs XXY/XY | K-S test | 8.5424E-09 |
| Fig. 5K | chrX lncRNA - XXXXY/XY vs XXY/XY | K-S test | 1.1366E-05 |
| Fig. 5K | chrX lncRNA - XXXY/XY vs XXXXY/XY | K-S test | 3.8536E-01 |
| Fig. 5L | autosome lncRNA - XX/XY vs XXY/XY | K-S test | 0.0000E+00 |
| Fig. 5L | autosome lncRNA - XXXY/XY vs XXY/XY | K-S test | 0.0000E+00 |
| Fig. 5L | autosome lncRNA - XXXXY/XY vs XXY/XY | K-S test | 0.0000E+00 |
| Fig. 5L | autosome lncRNA - XXXY/XY vs XXXXY/XY | K-S test | 3.5033E-10 |
| Fig. 6A | trisomy13 - cis-TF vs trans-TF | K-S test | 3.0743E-01 |
| Fig. 6C | trisomy18 - cis-TF vs trans-TF | K-S test | 2.3959E-02 |
| Fig. 6E | trisomy21 - cis-TF vs trans-TF | K-S test | 1.8422E-07 |
| Fig. 6A | trisomy13 cis-TF | normality test | 7.2979E-03 |
| Fig. 6A | trisomy13 trans-TF | normality test | 0.0000E+00 |
| Fig. 6C | trisomy18 cis-TF | normality test | 8.8798E-01 |
| Fig. 6C | trisomy18 trans-TF | normality test | 0.0000E+00 |
| Fig. 6E | trisomy21 cis-TF | normality test | 7.4278E-04 |
| Fig. 6E | trisomy21 trans-TF | normality test | 0.0000E+00 |
